# Neural networks to learn protein sequence-function relationships from deep mutational scanning data

**DOI:** 10.1101/2020.10.25.353946

**Authors:** Sam Gelman, Sarah A. Fahlberg, Pete Heinzelman, Philip A. Romero, Anthony Gitter

**Author notes:** These authors contributed equally to this work (Philip A. Romero, Anthony Gitter).

## Abstract

The mapping from protein sequence to function is highly complex, making it challenging to predict how sequence changes will affect a protein’s behavior and properties. We present a supervised deep learning framework to learn the sequence-function mapping from deep mutational scanning data and make predictions for new, uncharacterized sequence variants. We test multiple neural network architectures, including a graph convolutional network that incorporates protein structure, to explore how a network’s internal representation affects its ability to learn the sequence-function mapping. Our supervised learning approach displays superior performance over physics-based and unsupervised prediction methods. We find networks that capture nonlinear interactions and share parameters across sequence positions are important for learning the relationship between sequence and function. Further analysis of the trained models reveals the networks’ ability to learn biologically meaningful information about protein structure and mechanism. Finally, we demonstrate the models’ ability to navigate sequence space and design new proteins beyond the training set. We applied the GB1 models to design a sequence that binds to IgG with substantially higher affinity than wild-type GB1. Our software is available from https://github.com/gitter-lab/nn4dms.

**Significance:** Understanding the relationship between protein sequence and function is necessary to design new and useful proteins with applications in bioenergy, medicine, and agriculture. The mapping from sequence to function is tremendously complex because it involves thousands of molecular interactions that are coupled over multiple lengths and timescales. In this work, we show neural networks can learn the sequence-function mapping from large protein datasets. Neural networks are appealing for this task because they can learn complicated relationships from data, make few assumptions about the nature of the sequence-function relationship, and can learn general rules that apply across the length of the protein sequence. We demonstrate the learned models can be applied to design new proteins with properties that exceed natural sequences.

## Introduction

Understanding the mapping from protein sequence to function is important for describing natural evolutionary processes, diagnosing genetic disease, and designing new proteins with useful properties. This mapping is shaped by thousands of intricate molecular interactions, dynamic conformational ensembles, and nonlinear relationships between biophysical properties. These highly complex features make it challenging to model and predict how changes in amino acid sequence affect function.

The volume of protein data has exploded over the last decade with advances in DNA sequencing, three-dimensional structure determination, and high-throughput screening. With this increasing data, statistics and machine learning approaches have emerged as powerful methods to understand the complex mapping from protein sequence to function. Unsupervised learning methods such as EVmutation [1] and DeepSequence [2] are trained on large alignments of evolutionarily related protein sequences. These methods can model a protein family’s native function, but they are not capable of predicting specific protein properties that were not subject to long-term evolutionary selection. In contrast, supervised methods learn the mapping to a specific protein property directly from sequence-function examples. Many prior supervised learning approaches have limitations such as the inability to capture nonlinear interactions [3, 4], poor scalability to large datasets [5], making predictions only for single-mutation variants [6], or a lack of available code [7]. Other learning methods leverage multiple sequence alignments and databases of annotated genetic variants to make qualitative predictions about a mutation’s effect on organismal fitness or disease, rather than making quantitative predictions of molecular phenotype [8–10]. There is a current need for general, easy to use supervised learning methods that can leverage large sequence-function datasets to predict specific molecular phenotypes with the high accuracy required for protein design. We address this need with a usable software framework that can be readily adopted for new proteins [11].

We present a deep learning framework to learn protein sequence-function relationships from large-scale data generated by deep mutational scanning experiments. We train supervised neural networks to learn the mapping from sequence to function. These trained networks can then generalize to predict the functions of previously unseen sequences. We examine network architectures with different representational capabilities including linear regression, nonlinear fully connected networks, and convolutional networks that share parameters. Our supervised modeling approach displays strong predictive accuracy on five diverse deep mutational scanning datasets and compares favorably to state-of-the-art physics-based and unsupervised prediction methods. Across the different architectures tested, we find networks that capture nonlinear interactions and share information across sequence positions display the greatest predictive performance. We explore what our neural network models have learned about proteins and how they comprehend the sequence-function mapping. The convolutional neural networks learn a protein sequence representation that organizes sequences according to their structural and functional differences. In addition, the importance of input sequence features displays a strong correspondence to the protein’s three-dimensional structure and known key residues. Finally, we used an ensemble of the supervised learning models to design five GB1 sequences with varying distance from wild-type. We experimentally characterized these sequences and found the top design binds to IgG with at least an order of magnitude higher affinity than wild-type GB1.

## Results

### A deep learning framework to model the sequence-function mapping

Neural networks are capable of learning complex, nonlinear input-output mappings, extracting meaningful, higher-level features from raw inputs, and generalizing from training data to new, unseen inputs [12]. We develop a deep learning framework to learn from large-scale sequence-function data generated by deep mutational scanning. Deep mutational scanning data consists of thousands to millions of protein sequence variants that each have an associated score that quantifies their activity or fitness in a high-throughput function assay [13]. We encode the protein sequences with a featurization that captures the identity and physicochemical properties of each amino acid at each position. Our approach encodes the entire protein sequence and thus can represent multi-mutation variants. We train a neural network to map the encoded sequences to their associated functional scores. Once trained, the network generalizes and can predict functional scores for new, unseen protein variants (Fig. 1a).

**Figure 1.**
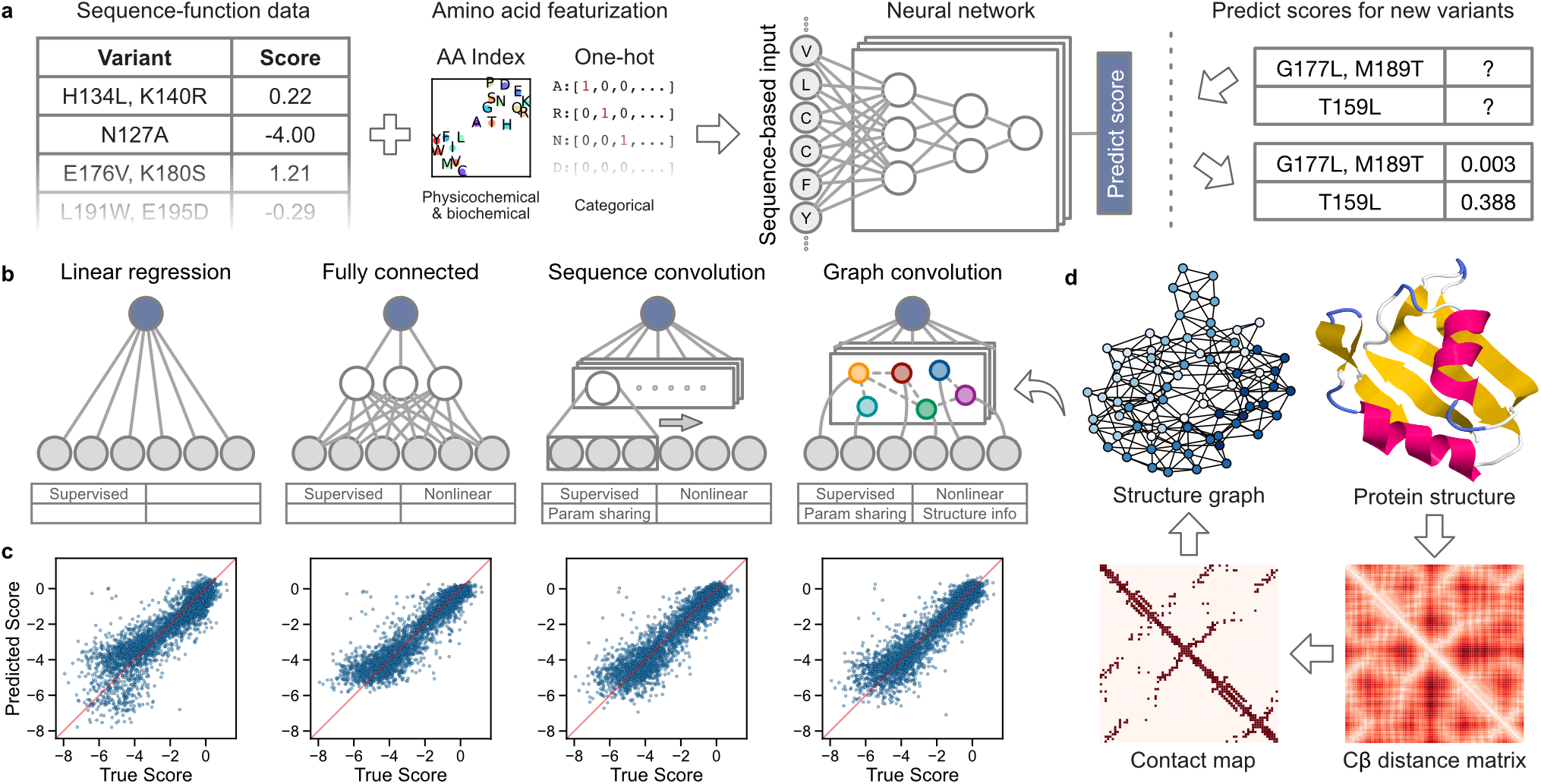
Overview of our supervised learning framework. (a) We use sequence-function data to train a neural network that can predict the functional score of protein variants. The sequence-based input captures physicochemical and biochemical properties of amino acids and supports multiple mutations per variant. The trained network can predict functional scores for previously uncharacterized variants. (b) We tested linear regression and three types of neural network architectures: fully connected, sequence convolutional, and graph convolutional. (c) Scatter plots showing performance of trained networks on the Pab1 dataset. (d) Process of generating the protein structure graph for Pab1. We create the protein structure graph by computing a residue distance matrix from the protein’s three-dimensional structure, thresholding the distances, and converting the resulting contact map to an undirected graph. The structure graph is the core part of the graph convolutional neural network.

We test four supervised learning models to explore how different internal representations influence the ability to learn the mapping from protein sequence to function: linear regression and fully connected, sequence convolutional, and graph convolutional neural networks (Fig. 1b). Linear regression serves as a simple baseline because it cannot capture dependencies between sites, and thus all residues make additive contributions to the predicted fitness. Fully connected networks incorporate multiple hidden layers and nonlinear activation functions, enabling them to learn complex nonlinearities in the sequence-to-function mapping. In contrast to linear regression, fully connected networks are capable of modeling how combinations of residues jointly affect function beyond simple additive effects. These non-additive effects are known as mutational epistasis [14, 15]. Neither linear regression nor fully connected networks are able to learn meaningful weights for amino acid substitutions that are not directly observed in the training set.

Convolutional neural networks have parameter sharing architectures that enable them to learn higher-level features that generalize across different sequence positions. They learn convolutional filters that identify patterns across different parts of the input. For example, a filter may learn to recognize the alternating pattern of polar and nonpolar amino acids commonly observed in beta strands. Applying this filter would enable the network to assess beta strand propensity across the entire input sequence and relate this higher-level information to the observed protein function. Importantly, the filter parameters are shared across all sequence positions, enabling convolutional networks to make meaningful predictions for mutations that were not directly observed during training. We develop a sequence-based convolutional network that integrates local sequence information by applying filters using a sliding window across the amino acid sequence. We also develop a structure-based graph convolutional network that integrates three-dimensional structural information and may allow the network to learn filters that correspond to structural motifs. The graph convolutional network applies filters to neighboring nodes in a graph representation of the protein’s structure. The protein structure graph consists of a node for each residue and an edge between nodes if the residues are within a specified distance in three-dimensional space (Fig. 1d).

### Evaluating models learned from deep mutational scanning data

We evaluated the predictive performance of the different network architectures on five diverse deep mutational scanning datasets representing proteins of varying sizes, folds, and functions: avGFP, Bgl3, GB1, Pab1, and Ube4b (Table 1and Fig. 2a). These datasets range in size from ~25,000 to ~500,000 sequence-score examples. We randomly split each dataset into training, tuning, and testing sets to optimize hyperparameters and evaluate predictive performance on data that was not seen during training. The learned models displayed excellent test set predictions for most datasets, with Pearson’s correlation coefficients ranging from 0.55 to 0.98 (Fig. 2b). The trends are generally similar using Spearman’s correlation coefficient (Fig. S1), although the differences between linear regression and the neural networks are smaller.

**Table 1.** Deep mutational scanning datasets. We evaluated the models on deep mutational scanning datasets representing proteins of varying sizes, folds, and functions.

|  | Description | Organism | Molecular function | Selection | Length | Variants | Ref |
| --- | --- | --- | --- | --- | --- | --- | --- |
| avGFP | Green fluorescent protein | A. victoria | Fluorescence | Brightness | 237 | 54,024 | [16] |
| Bgl3 | Beta glucosidase | Streptococcus sp. | Hydrolysis of β-glucosidic linkages | Enzymatic activity | 501 | 26,653 | [17] |
| GB1 | IgG-binding domain of protein G | Streptococcus sp. | IgG-binding | IgG-Fc binding | 56 | 536,084 | [15] |
| Pab1 | RRM domain of Pab1 | S. cerevisiae | poly(A)-binding | mRNA binding | 75 | 40,852 | [18] |
| Ube4b | U-box domain of E4B | M. musculus | Ubiquitin activating enzyme activity | Ubiquitin ligase activity | 102 | 98,297 | [19] |

**Figure 2.**
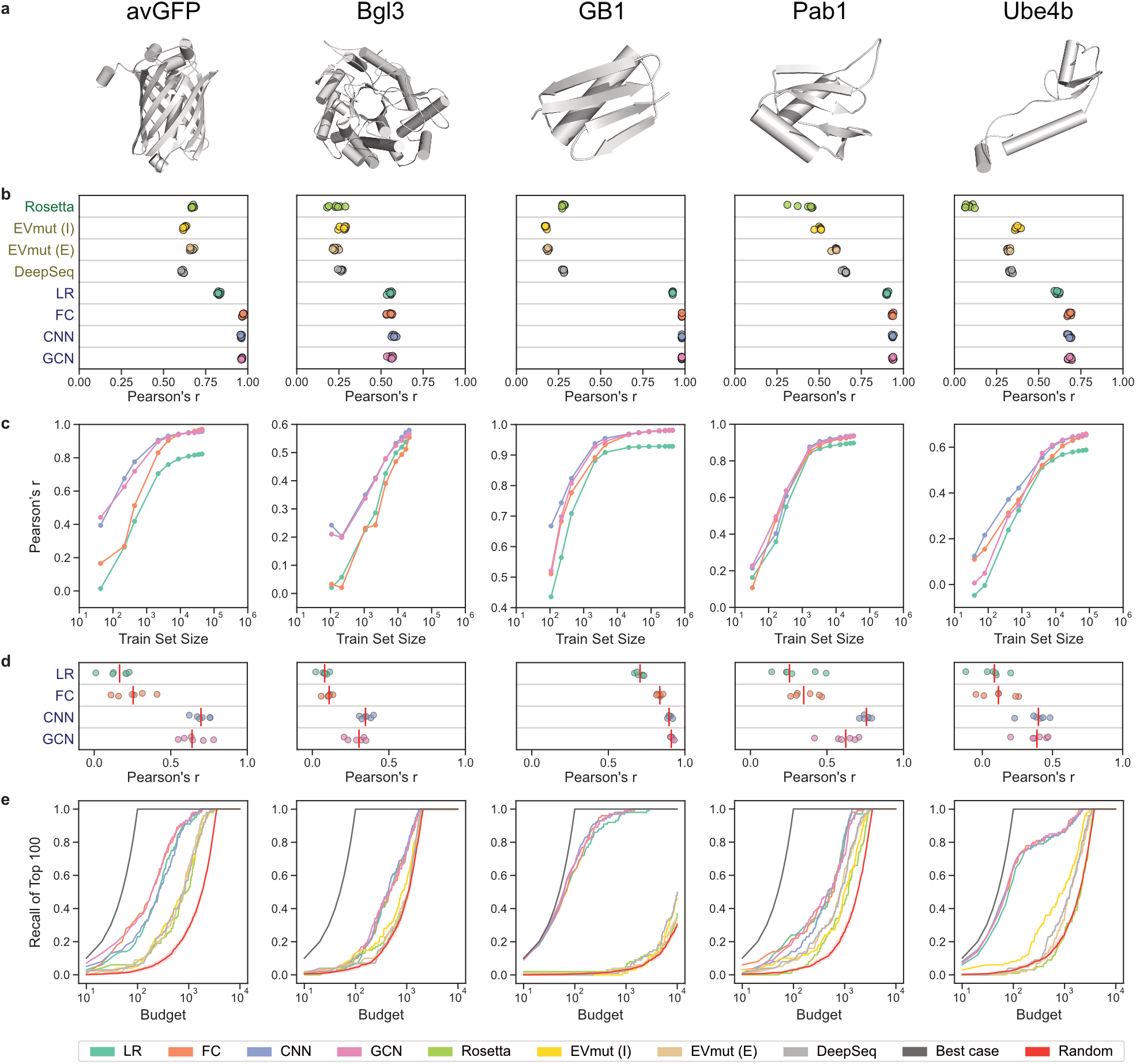
Evaluation of neural networks and comparison to unsupervised methods. (a) Three-dimensional protein structures. (b) Pearson’s correlation coefficient between true and predicted scores for Rosetta, EVmutation, DeepSequence, linear regression (LR), fully connected network (FC), sequence convolutional network (CNN), and graph convolutional network (GCN). EVmutation (I) refers to the independent formulation of the model that does not include pairwise interactions. EVmutation (E) refers to the epistatic formulation of the model that does include pairwise interactions. Each point corresponds to one of seven random train-tune-test splits. (c) Correlation performance of supervised models trained with reduced training set sizes. (d) Model performance when making predictions for variants containing mutations that were not seen during training (mutational extrapolation). Each point corresponds to one of six replicates, and the red vertical line denotes the median. (e) The fraction of the true 100 best-scoring variants identified by each model’s ranking of variants with the given budget. The random baseline is shown with the mean and a 95% confidence interval.

For comparison, we also evaluated the predictive performance of established physics-based and unsupervised learning methods Rosetta [20], EVmutation [1], and DeepSequence [2], which are not trained using the deep mutational scanning data. Our supervised learning approach achieves superior performance to these other methods on all five protein datasets, demonstrating the benefit of training directly on sequence-function data (Fig. 2b). This result is unsurprising because supervised models are tailored to the specific protein property and sequence distribution in the dataset. Rosetta predictions consider the energetics of the protein structure and therefore do not capture the more specific aspects of protein function. Unsupervised methods such as EVmutation and DeepSequence are trained on natural sequences and thus only capture aspects of protein function directly related to natural evolution. Despite their lower performance, physics-based and unsupervised methods have the benefit of not requiring large-scale sequence-function data, which is often difficult and expensive to acquire.

The different supervised models displayed notable trends in predictive performance across the datasets. The nonlinear models outperformed linear regression, especially on variants with low scores (Fig. S2), high epistasis (Fig. S3), and in the case of avGFP, larger numbers of mutations (Fig. S4). The three nonlinear models performed similarly when trained and evaluated on the full training set and testing sets. However, the convolutional networks achieved a better mean squared error when evaluating single-mutation variants in Pab1 and GB1 (Fig. S4). For most proteins, the convolutional networks also had superior performance when trained on smaller training sets (Fig. 2c).

The quantitative evaluations described thus far involve test set variants that have similar characteristics to the training data. We also tested the ability of the models to extrapolate to more challenging test sets. In mutational extrapolation, the model makes predictions for variants containing mutations that were not seen during training. The model must generalize based on mutations that may occur in the same or other positions. The convolutional networks achieved strong performance for one dataset (r > 0.9), moderate performance for two additional datasets (r > 0.6), and outperformed linear regression and fully connected networks across all datasets (Fig. 2d). In positional extrapolation, the model makes predictions for variants containing mutations in positions that were never modified in the training data. The performance of all models is drastically reduced (Fig. S5), highlighting the difficulty of this task [21]. In theory, the parameter sharing inherent to convolutional networks allows them to generalize the effects of mutations across sequence positions. This capability may explain the convolutional networks’ superior performance with reduced training set sizes and mutational extrapolation. However, it is still difficult for the convolutional networks to perform well when there are no training examples of mutations in a particular position, such as in positional extrapolation.

The sequence convolutional and graph convolutional networks displayed similar performance across all evaluation metrics, despite the inclusion of three-dimensional protein structure information in the graph topology. To assess the impact of integrating protein structure in the neural network architecture, we created graph convolutional networks with misspecified baseline graphs that were unrelated to the protein structure. These baseline graphs include shuffled, disconnected, sequential, and complete graph structures (Fig. S6). We found networks trained using these misspecified baseline graphs had accuracy similar to networks trained with the actual protein structure graph, indicating that protein structure information is contributing little to the model’s performance (Fig. S7). We also trained the convolutional networks with and without a final fully connected layer and found this fully connected layer was more important than a correctly specified graph structure. In almost all cases, this final fully connected layer helps overcome the misspecified graph structure (Fig. S7). Overall, these results suggest the specific convolutional window is not as critical as sharing parameters across different sequence positions and integrating information with a fully connected layer.

The goal of protein engineering is to identify optimized proteins, and models can facilitate this process by predicting high activity sequences from an untested pool of sequences. Pearson’s correlation captures a model’s performance across all variants, but it does not provide information regarding a model’s ability to retrieve and rank high scoring variants. We evaluated each model’s ability to predict the highest scoring variants within a given experimental testing budget (Fig. 2e). We calculated recall by asking each model to identify the top N variants from the test set, where N is the budget, and evaluating what fraction of the true top 100 variants were covered in this predicted set. The supervised models consistently achieve higher recall than Rosetta and the unsupervised methods, although the differences are small for Pab1 and Bgl3. In practice, the budget depends on the experimental costs of synthesizing and evaluating variants of the given protein. For GB1, a feasible budget may be 100 variants, and the supervised models can recall over 60% of the top 100 sequences with that budget.

Another important performance metric for protein engineering is the ability to prioritize variants that have greater activity than the wild-type protein. We calculated the mean and max scores of the top N predicted test set variants ranked by each model (Figs. S8 and S9). We find the variants prioritized by the supervised models have greater functional scores than wild-type on average, even when considering variants ranked beyond the top thousand sequences for some datasets. In contrast, Rosetta and the unsupervised models generally prioritize variants with mean scores worse than wild-type. The max score of the prioritized variants is also important because it represents the best variant suggested by the model. We find nearly all models are able to prioritize variants with a max score greater than wild-type. The relative performance of each model is dependent on the dataset. Notably, the unsupervised methods perform very well on Bgl3, with EVmutation identifying the top variant with a budget of 20. Meanwhile, the supervised methods perform very well on Ube4b, prioritizing a variant with the true max score with a budget as small as 5 variants.

### Role of data quality in learning accurate sequence-function models

The performance of the supervised models varied substantially across the five protein datasets. For example, the Pearson correlation for the Bgl3 models was ~0.4 lower than the GB1 models. Although it is possible some proteins and protein families are intrinsically more difficult to model, practical considerations such as the size and quality of the deep mutational scanning dataset could also affect protein-specific performance. Deep mutational scanning experiments use a high-throughput assay to screen an initial gene library and isolate variants with a desired functional property. The initial library and the isolated variants are sequenced, and a fitness score is computed for each variant based on the frequency of reads in both sets. The quality of the calculated fitness scores depends on the sensitivity and specificity of the high-throughput assay, the number of times each variant was characterized in the high-throughput assay, and the number of DNA sequencing reads per variant. If any one of these factors is too low, the resulting fitness scores will not reflect the true fitness values of the characterized proteins, which will make it more difficult for a model to learn the underlying sequence to function mapping.

We assessed how experimental factors influence the success of supervised learning by resampling the full GB1 dataset to generate simulated datasets with varying protein library sizes and numbers of DNA sequencing reads. The library size is the number of unique variants screened in the deep mutational scan. The GB1 dataset is ideal for this analysis because it contains most single and double mutants and has a large number of sequencing reads per variant. We trained sequence convolutional models on each simulated dataset and tested each network’s predictions on a “true”, non-resampled test set (Fig. 3). Models trained on simulated datasets with small library sizes performed poorly because there were not sufficient examples to learn the sequence-function mapping. This result is expected and is in line with the performance of models trained on reduced training set sizes on the original GB1 dataset (Fig. 2c). Interestingly, we also found datasets with large library sizes can perform poorly if there are not sufficient DNA sequencing reads to reliably estimate the frequency of each variant. This highlights a trade-off between the number of sequence-function examples in a dataset and the quality of its fitness scores. Given a fixed sequencing budget, there exists an optimal intermediate library size that balances these two competing factors. The Bgl3 dataset’s poor performance may be the result of having too many unique variants without sufficient sequencing coverage, resulting in a low number of reads per variant and therefore unreliable fitness scores. Future deep mutational scanning libraries could be designed to maximize their size and diversity while ensuring that each variant will have sufficient reads within sequencing throughput constraints.

**Figure 3.**
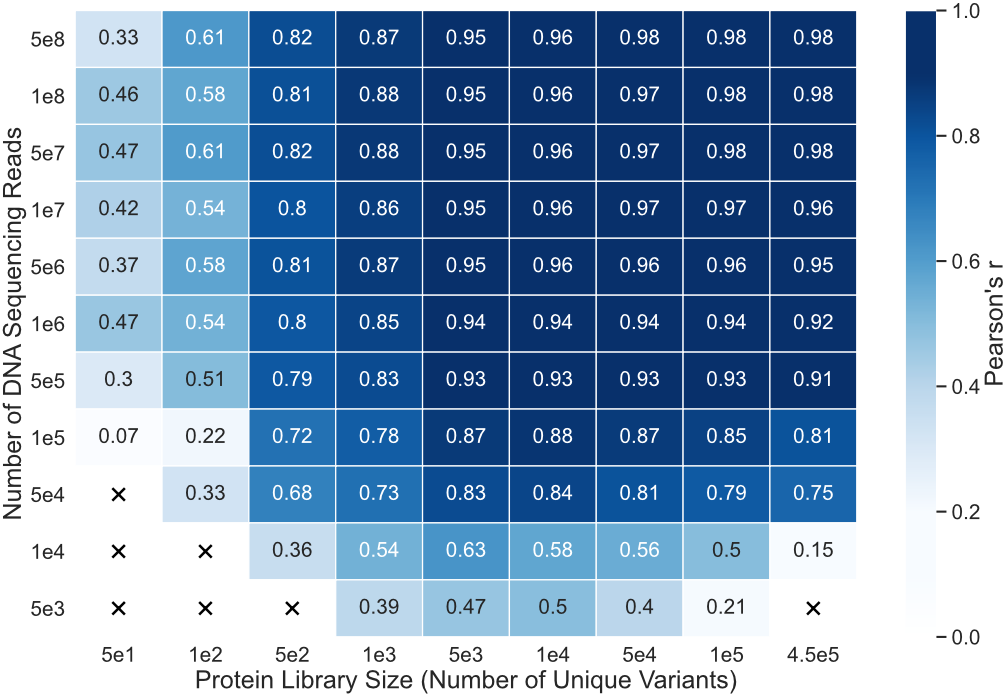
Trade-off between library size and number of sequencing reads. Performance of sequence convolutional models trained on GB1 datasets that have been resampled to simulate different combinations of protein library size and number of sequencing reads in the deep mutational scan. An “X” signifies the combination of library size and number of reads produced a dataset with fewer than 25 variants and was therefore excluded from the experiment. Having a large library size can be detrimental to supervised model performance if there are not enough reads to calculate reliable functional scores.

### Learned models provide insight into protein structure and mechanism

Our neural networks transform the original amino acid features through multiple layers to map to an output fitness value. Each successive layer of the network constructs new latent representations of the sequences that capture important aspects of protein function. We can visualize the relationships between sequences in these latent spaces to reveal how the networks learn and comprehend protein function. We used Uniform Manifold Approximation and Projection (UMAP) [22] to visualize test set sequences in the latent space at the last layer of the GB1 sequence convolutional network (Fig. 4a). The latent space organizes the test set sequences based on their functional score, demonstrating that the network’s internal representation, which was learned to predict function of the training set examples, also generalizes to capture the sequence-function relationship of the new sequences. The latent space features three prominent clusters of low-scoring variants that may correspond to different mechanisms of disrupting GB1 function. Two clusters, referred to as “G1” and “G2”, contain variants with mutations in core residues near the protein’s N and C termini, respectively (Fig. S10). Mutations at these residues may disrupt the protein’s structural stability and thus decrease the activity measured in the deep mutational scanning experiment [15]. Residue cluster “G3” contains variants with mutations at the IgG-binding interface, and these likely decrease activity by disrupting key binding interactions. This clustering of variants based on different molecular mechanisms suggests the network is learning biologically meaningful aspects of protein function.

**Figure 4.**
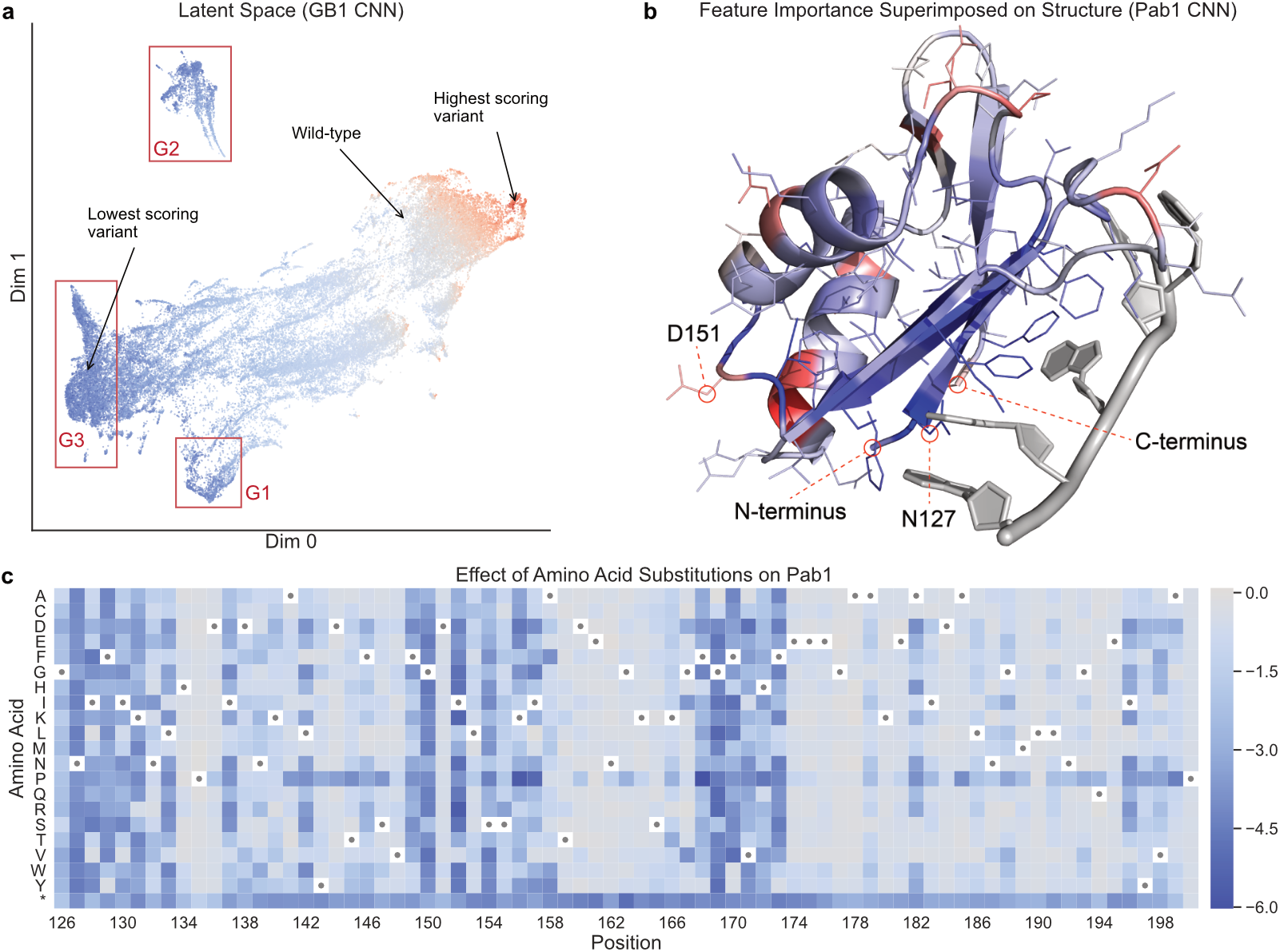
Neural network interpretation. (a) A UMAP projection of the latent space of the GB1 sequence convolutional network, as captured at the last internal layer of the network. In this latent space, similar variants are grouped together based on the transformation applied by the network to predict the functional score. Variants are colored by their true functional score, where red represents high-scoring variants and blue represents low-scoring variants. The clusters marked G1 and G2 correspond to variants with mutations at core residues near the start and end of the sequence, respectively. Cluster G3 corresponds to variants with mutations at surface interface residues. (b) Integrated gradients feature importance values for the Pab1 sequence convolutional network, aggregated at each sequence position and superimposed on the protein’s three-dimensional structure. Blue represents positions with negative attributions, meaning mutations in those positions push the network to output lower scores, and red represents positions with positive attributions. (c) A heat map showing predictions for all single mutations from the Pab1 sequence convolutional network. Wild-type residues are indicated with dots, and * is the stop codon. Most single mutations are predicted to be neutral or deleterious.

We can also use the neural network models to understand which sequence positions have the greatest influence on protein function. We computed integrated gradients attributions [23] for all training set variants in Pab1 and mapped these values onto the three-dimensional structure (Fig. 4b). Pab1’s sequence positions display a range of attributions spanning from negative to positive, where a negative attribution indicates mutations at that position decrease the protein’s activity. Residues at the RNA-binding interface tend to display negative attributions, with the key interface residue N127 having the largest negative attribution. The original deep mutational scanning study found residue N127 cannot be replaced with any other amino acid without significantly decreasing Pab1 binding activity [18]. Position D151 has one of the largest positive attributions, which is consistent with the observation that aspartic acid (D) is uncommon at position 151 in naturally occurring Pab1 sequences [18]. The sequence convolutional network is able to learn biologically relevant information directly from raw sequence-function data, without the need to specify detailed molecular mechanisms.

Finally, we used the Pab1 sequence convolutional network to make predictions for all possible single-mutation variants (Fig. 4c). The resulting heat map highlights regions of the Pab1 sequence that are intolerant to mutations and shows that mutations to proline are deleterious across most sequence positions. It also demonstrates the network’s ability to predict scores for amino acids that were not directly observed in the dataset. The original deep mutational scan characterized 1,244 single-mutation variants, yet the model can make predictions for all 1,500 possible single-mutation variants. For example, mutation F170P was not experimentally observed, but the model predicts it will be deleterious because proline substitutions at other positions are often highly deleterious. This generalization to amino acid substitutions not observed in the data is only possible with models that share parameters across sequence positions.

### Designing distant protein sequences with learned models

Our trained neural networks describe the mapping from sequence to function for a given protein family. These models can be used to design new sequences that were not observed in the original deep mutational scanning dataset and may have improved function. The protein design process involves extrapolating a model’s predictions to distant regions of sequence space. Because the models were trained and evaluated only on sequences with local changes with respect to the wild-type, it is unclear how these out-of-distribution predictions will perform.

We tested the ability of our supervised models to generalize beyond the training data by designing a panel of new GB1 variants with varying distances from the wild-type sequence (Fig. 5a). GB1 is a small 8 kDa domain from streptococcal protein G that binds to the Fc domain of mammalian immunoglobulin G (IgG). GB1’s structure is composed of one alpha-helix that is packed into a four-stranded beta-sheet. GB1’s interaction with IgG is largely mediated by residues in the alpha-helix and third beta-strand.

**Figure 5.**
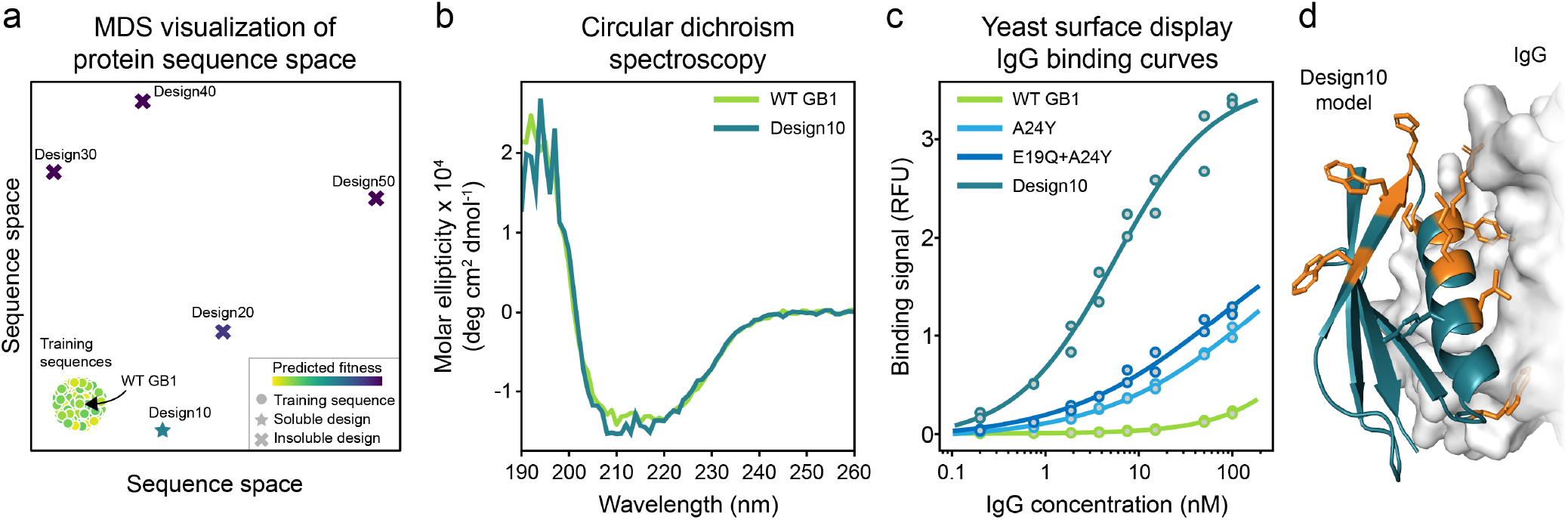
Neural network-based protein design. (a) Multidimensional scaling (MDS) sequence space visualization of the GB1 training sequences and the five designed proteins. Design10 - Design50 are progressively further from the training distribution. Design10 expressed as a soluble protein, while the more distant designs were insoluble. (b) Circular dichroism spectra of purified wild-type GB1 and Design10. Both proteins display highly similar spectra that are indicative of alpha helical protein structures. (c) IgG binding curves of wild-type GB1 variants. Design10 displays substantially higher binding affinity than wild-type GB1, A24Y, and E19Q+A24Y. All measurements were performed in duplicate. (d) The locations of Design10’s ten mutations (shown in orange) relative to the IgG binding interface. The Design10 structure was predicted *de novo* using Rosetta.

The design process was guided by an ensemble of the four models (linear regression and fully connected, sequence convolutional, and graph convolutional networks) to fully leverage different aspects of the sequence-function mapping captured by each model. We used a random-restart hill climbing algorithm to search over GB1 sequence space for designs that maximize the minimum predicted fitness over the four models. Maximizing the minimum predicted fitness over the four models ensures that every model predicts the designed sequences to have high fitness. We applied this sequence optimization method to design five GB1 variants with increasing numbers of mutations (10, 20, 30, 40, and 50) from wild-type, representing sequence identities spanning from 82% to 11% (Table S1). We expect designs with fewer mutations to be more likely to fold and function because they are more similar to the training data.

We experimentally tested the five GB1 designs by synthesizing their corresponding genes and expressing them in E. coli. We found the 10-mutant design, referred to as Design10, was expressed as a soluble protein, but the more distant designs were insoluble (Fig. 5a). We were unable to further characterize Designs 20-50 because their insoluble expression prevented downstream protein purification. We performed circular dichroism spectroscopy on Design10 and found it had a nearly identical spectra to wild-type GB1 suggesting they have similar secondary structure content (Fig. 5b).

The original GB1 deep mutational scan measured binding to the Fc region of IgG; therefore, our supervised models should capture a variant’s binding affinity. We tested Design10’s ability to bind to IgG using a yeast display binding assay. We also tested wild-type GB1 and the top single (A24Y) and double (E19Q+A24Y) mutants from the original deep mutational scanning dataset. We found Design10 binds to IgG with a *K_d_* of 5 nM, which is substantially higher affinity than wild-type GB1, A24Y, or E19Q+A24Y (Fig. 5c). We were unable to precisely determine wild-type GB1, A24Y, or E19Q+A24Y’s dissociation constants because our assay could not reliably measure binding at IgG concentrations above 100 nM. The data showed qualitative trends where WT had the lowest affinity, followed by A24Y, and then E19Q+A24Y. Our measurements indicate that wild-type GB1’s Kd is well above 100 nM, which is consistent with measurements from the literature that have found this interaction is in the 250-900 nM range [24, 25]. Based on our estimates and others’ previous measurements, we conservatively estimate that Design10 binds human IgG with at least 20-fold higher affinity than wild-type GB1.

Closer inspection of the Design10 sequence revealed that it was not simply composed of the top ten single mutations for enrichment and even included four mutations whose individual effects on predicted the functional score ranked below the top 300. In addition, Design10’s predicted score was more than two times greater than the variant comprising the top ten single mutations. This highlights the ability of the design process to capture nonlinear interactions and leverage synergies between sites. We also evaluated the robustness of our findings by re-running the 10-mutant design process 100 independent times and evaluating the diversity of the designs (Table S2).

We built a model of Design10’s three-dimensional structure using Rosetta *de novo* structure prediction (Fig. 5d). Design10’s predicted structure aligns to the wild-type GB1 crystal structure with 0.9Å C*α* root-mean-square deviation. Design10’s actual structure is likely very similar to wild-type GB1 given their high sequence identity, similar circular dichroism spectra, and the small deviation between Design10’s *de novo* predicted structure and the experimental GB1 structure. Inspection of Design10’s predicted structure revealed that many of its mutations were concentrated near the IgG binding interface, and this may help to explain its large increase in IgG binding affinity. We also evaluated Rosetta models for Designs 20-50 and found no obvious reasons why they might have failed to express.

## Discussion

We have presented a supervised learning framework to infer the protein sequence-function mapping from deep mutational scanning data. Our supervised models work best when trained with large-scale datasets, but they can still outperform physics-based and unsupervised prediction methods when trained with only hundreds of sequence-function examples. Unsupervised methods remain appealing for proteins with very little or no sequence-function data available. Among the supervised models, linear regression displayed the lowest performance due to its inability to represent interactions between multiple mutations. Despite that limitation, linear regression still performed fairly well because mutations often combine in an additive manner [26]. The convolutional networks outperformed linear regression and fully connected networks when trained with fewer training examples and when performing mutational extrapolation. The parameter sharing inherent to convolutional networks can improve performance by allowing generalization of the effects of mutations across different sequence positions. However, in the five datasets we tested, even the convolutional networks could not accurately generalize when entire sequence positions were excluded from the training data. It was that surprising graph convolutions that incorporate protein structure did not improve performance over sequence-based convolutions. The comparable performance could be the result of the networks’ ability to compensate with fully connected layers, the lack of sequence diversity in the deep mutational scanning data, or the specific type of graph neural network architecture used. We are unable to determine which of these factors had the greatest influence.

Our analysis of how data quality influences the ability to learn the sequence-function mapping can be considered when designing future deep mutational scanning experiments. We found a model’s predictive performance is determined not only by the number of sequence-function training examples but also by the quality of the estimated functional scores. Therefore, in a deep mutational scanning experiment it may be preferable to limit the total number of unique variants analyzed to ensure that each variant has sufficient sequencing reads to calculate accurate functional scores. Any missing mutations can then be imputed with a convolutional network to overcome the smaller dataset size.

Recent studies have examined supervised learning methods capable of scaling to deep mutational scanning datasets. One study benchmarked combinations of supervised learning methods and protein sequence encodings [7]. Consistent with our results, it found sequence convolutional neural networks with amino acid property-based features tended to perform better than alternatives. Some algorithms specialize in modeling epistasis. Epistatic Net [27] introduced a neural network regularization strategy to limit the number of epistatic interactions. Other approaches focused on the global epistasis that arises due to a nonlinear transformation from a latent phenotype to the experimentally-characterized function [28, 29]. Protein engineering with UniRep [30] showed that general global protein representations can support training function-specific supervised models with relatively few sequence-function examples. ECNet pioneered an approach for combining global protein representations, local information about residue co-evolution, and protein sequence features [31]. Across tens of deep mutational scanning datasets, ECNet was almost always superior to unsupervised learning models and models based only on a global protein representation. Future work can explore how to best combine global protein representations, local residue co-evolution features, and graph encodings of protein structure to learn predictive models for specific protein functions, including for proteins that have little experimental data available. Despite their similar performance to sequence convolutional networks in our study, graph convolutional networks that integrate three-dimensional structural information remain enticing because of successes on other protein modeling tasks [32–35] and rapid developments in graph neural network architectures [36].

Another challenging future direction will be assessing how well trained models extrapolate to sequences with higher-order mutations [21, 37]. As a proof-of-concept, we designed distant GB1 sequences with 10s of mutations from wild-type. The 10-mutant design (Design10) had substantially stronger IgG binding affinity than wild-type GB1, but the four sequences with more mutations did not express as soluble proteins. The tremendous success of Design10 is encouraging considering how few designed sequences we tested and the many opportunities to improve upon our limited exploration of model-guided design. The model predictions can be improved through more sophisticated ensembling and uncertainty estimation. Our hill climbing sequence optimization strategy can be replaced by specialized methods that allow supervised models to efficiently explore new parts of a sequence space [38–41].

Machine learning is revolutionizing our ability to model and predict the complex relationships between protein sequence, structure, and function [42, 43]. Supervised models of protein function are currently limited by the availability and quality of experimental data but will become increasingly accurate and general as researchers continue to experimentally characterize protein sequence space [44]. Other important machine learning advances relevant to protein engineering include generative modeling to sample non-natural protein sequences [34, 45, 46], language models to learn protein representations from diverse natural sequences [47–50], and strategies to incorporate machine learning predictions into directed evolution experiments [51–53]. These approaches could make possible a new generation of data-driven protein engineering.

## Methods

### Datasets

We tested our supervised learning approach on five deep mutational scanning datasets: avGFP [16], Bgl3 [17], GB1 [15], Pab1 [18], and Ube4b [19]. We selected these publicly available datasets because they correspond to diverse proteins and contain variants with multiple amino acid substitutions. The avGFP, Pab1, and Ube4b datasets were published with precomputed functional scores, which we used directly as the target scores for our method. For GB1 and Bgl3, we computed functional scores from raw sequencing read counts using Enrich2 [54]. We filtered out variants with fewer than five sequencing reads and ran Enrich2 using the “Log Ratios (Enrich2)” scoring method and the “Wild Type” normalization method. See Table 1 for additional details about the datasets.

### Protein sequence encoding

We encoded each variant’s amino acid sequence using a sequence-level encoding that supports multiple substitutions per variant. Each amino acid is featurized with its own feature vector, and the full encoded variant consists of the concatenated amino acid feature vectors. We featurize each amino acid using a two-part encoding made up of a one-hot encoding and an AAindex encoding. One-hot encoding captures the specific amino acid at each position. It consists of a length 21 vector where each position represents one of the possible amino acids or the stop codon. All positions are zero except the position of the amino acid being encoded, which is set to a value of one. AAindex encoding captures physicochemical and biochemical properties of amino acids from the AAindex database [55]. These properties include simple attributes, such as hydrophobicity and polarity, as well as more complex characteristics, such as average non-bonded energy per atom and optimized propensity to form a reverse turn. In total, there are 566 such properties that were taken from literature. These properties are partially redundant because they are aggregated from different sources. Therefore, we used principle component analysis to reduce the dimensionality to a length 19 vector, capturing 100% of the variance. We concatenated the one-hot and AAindex encodings to form the final representation for each amino acid. One benefit of this encoding is that it enables the use of convolutional networks, which leverage the spatial locality of the raw inputs to learn higher-level features via filters. Other types of encodings that do not have a feature vector for each residue, such as those that embed full amino acid sequences into fixed-size vectors, would not be as appropriate for convolutional networks because they do not have locality in the input that can be exploited by convolutional filters.

### Convolutional neural networks

We tested two types of convolutional neural networks: sequence convolutional and graph convolutional. These networks extract higher level features from the raw inputs using convolutional filters. Convolutional filters are sets of learned weights that identify patterns in the input and are applied across different parts of the input. The filters can output higher or lower values depending on whether the given input matches the pattern that the filters have learned to identify. We implemented a sequence convolutional network where the input is a one-dimensional amino acid sequence. The network applies filters using a sliding window across the input sequence, integrating information from amino acid sequence neighbors. The network applies filters at all valid sequence positions and does not pad the ends of the sequence with zeros.

Graph convolutional neural networks are similar to traditional convolutional networks, except graph convolutional networks operate on arbitrary graph structures rather than linear sequences or two-dimensional grids. Graph filters still capture spatial relationships in the input data, but those relationships are determined by neighboring nodes in the graph rather than neighboring characters in a sequence or neighboring pixels in a two-dimensional grid. In our case, we use a graph derived from the protein’s wild-type three-dimensional structure. This allows the network to more easily learn features that correspond to patterns of amino acid residues that are nearby in physical space.

We use the order-independent graph convolution operator described by Fout et al. [32]. It is considered order-independent because it does not impose an ordering on neighbor nodes. In an order-dependent formulation, different neighbor nodes would have different weights, but in the order-independent formulation, all neighbor nodes are treated identically and share the same weights. Each filter consists of a weight vector for the center node and a weight vector for the neighbor nodes that is shared among the neighbor nodes. For a set of filters, the output *z_i_* at a center node *i* is calculated using Equation 1, where *W_C_* is the center node’s weight matrix, *W_N_* is the neighbor nodes’ weight matrix, and *b* is the vector of biases, one for each filter. Additionally, *x_i_*, is the feature vector at node *i*, *N_i_* is the set of neighbors of node *i*, and σ is the activation function.

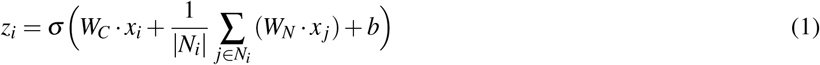

In this formulation, a graph consisting of nodes and edges is incorporated into each convolutional layer. Input features are placed at the graph’s nodes in the first layer. Outputs are computed at the node-level using input features from a given center node and corresponding neighbor nodes. Because output is computed for each node, graph structure is preserved between subsequent graph layers. The incoming signal from neighbor nodes is averaged to account for the variable numbers of neighbors. The window size of the filter is limited to the immediate neighbors of the current center node. Information from more distant nodes is incorporated through multiple graph convolutional layers. The final output of the network is computed at the graph-level with a single function score prediction for the entire graph.

### Protein structure as a graph

We encoded each protein’s wild-type structure as a graph and incorporated it into the architecture of the graph convolutional neural network (Fig. 1d). The protein structure graph is an undirected graph with a node for each amino acid residue and an edge between nodes if the residues are within a specified distance threshold in three-dimensional space. The distance threshold is a hyperparameter with a range of 4Å-10Å and was selected independently for each dataset during the hyperparameter optimization. We measure distances between residues via distances of the beta carbon atoms (C*β*) in angstroms. The protein structure graph for GB1 is based on Protein Data Bank (PDB) structure 2QMT. The protein structure graphs for the other four proteins are based on structures derived from Rosetta comparative modeling, using the RosettaCM protocol [56] with the default options. For the comparative modeling, we selected template structures from PDB that most closely matched the reference sequence of the deep mutational scanning data. In addition to the standard graph based on the protein’s structure, we tested four baseline graphs: a shuffled graph based on the standard graph but with shuffled node labels, a disconnected graph with no edges, a sequential graph containing only edges between sequential residues, and a complete graph containing all possible edges (Fig. S6). We used NetworkX [57] v2.3 to generate all protein structure and baseline graphs.

### Complete model architectures

We implemented linear regression and three types of neural network architectures: fully connected, sequence convolutional, and graph convolutional. Linear regression is implemented as a fully connected neural network with no hidden layers. It has a single output node that is fully connected to all input nodes. The other networks all have multiple layers. The fully connected network consists of some number of fully connected layers, and each fully connected layer is followed by a dropout layer with a 20% dropout probability. Finally, there is a single output node. The sequence and graph convolutional networks consist of some number of convolutional layers, a single fully connected layer with 100 hidden nodes, a dropout layer with a 20% dropout probability, and a single output node. We also trained sequence and graph convolutional networks without the fully connected layer or dropout layer for the analyses in Figure S7. We used the leaky rectified linear unit (LReLU) as the activation function for all hidden layers. A hyperparameter sweep determined the other key aspects of the model architectures, such as the number of layers, the number of filters, and the kernel size of filters (Fig. S11). We used Python v3.6 and TensorFlow [58] v1.14 to implement the models.

### Model training

We trained the networks using the Adam optimizer and mean squared error loss. We set the Adam hyperparameters to the defaults except for the learning rate and batch size, which were selected using hyperparameter sweeps. We used early stopping for all model training with a patience of 15 epochs and a minimum delta (the minimum amount by which loss must decrease to be considered an improvement) of 0.00001. We set the maximum possible number of epochs to 300. The original implementation over-weighted the last examples in an epoch when calculating the tuning set loss. This could have affected early stopping but had little to no effect in practice. We trained the networks on graphics processing units (GPUs) available at the University of Wisconsin-Madison via the Center for High Throughput Computing and the workload management system HTCondor [59]. We also used GPU resources from Argonne National Laboratory’s Cooley cluster. The GPUs we used included Nvidia GeForce GTX 1080 Ti, GeForce RTX 2080 Ti, and Tesla K80.

### Main experiment setup

We split each dataset into random training, tuning, and testing sets. The tuning set is sometimes referred to as the validation set and is used for hyperparameter optimization. This allowed us to train the models, tune hyperparameters, and evaluate performance on separate, non-overlapping sets of data. The training set was 81% of the data, the tuning set was 9%, and the testing set was 10%. This strategy supports the objective of training and evaluating models that fully leverage all available sequence-function data and make predictions for variants that have characteristics similar to the training data. There are other valid strategies that more directly test the ability of a model to generalize to mutations or positions that were not present in the training data, which we describe below.

We performed a hyperparameter grid search for each dataset using all possible combinations of the hyperparameters in Figure S11. The hyperparameters selected for one dataset did not influence the hyperparameters selected for any other dataset. For each type of supervised model (linear regression, fully connected, sequence convolutional, and graph convolutional), we selected the set of hyperparameters that resulted in the smallest mean squared error on the tuning set. The selected hyperparameters are listed in Table S3, and the number of trainable parameters in each selected model is listed in Table S4.

This is the main set of hyperparameters. For any subsequently trained models, such as those with reduced training set sizes, we performed smaller hyperparameter sweeps to select a learning rate and batch size, but all other hyperparameters that specify the network architecture were set to those selected in the main run.

To assess the robustness of the original train-tune-test splits, we created six additional random splits for each dataset. We tuned the learning rate and batch size independently for each split, however, the network architectures were fixed to those selected using the original split. There was a risk of overestimating performance on the new splits because the data used to tune the architectures from the original split may be present in the test sets of the new splits. However, the results showed no evidence of this type of overfitting. We report the performance on these new splits in Figures 2b and S1a. All other experiments used the original train-tune-test split.

For the random baseline in Figures S8 and S9, we generated 1,000 random rankings of the entire test set. Then for each ranking threshold N, we selected the first N variants from each ranking as the prioritized variants. We computed the mean (Fig. S8) and the max (Fig. S9) of each random ranking’s prioritized variants. Finally, we show the 95% confidence interval calculated as ± 1.96 times the standard deviation.

### Reduced training size setup

For the reduced training size experiment, we used the same 9% tuning and 10% testing sets as the main experiment. The reduced training set sizes were determined by percentages of the original 81% training pool. For each reduced size, we sampled five random training sets of the desired size from the 81% training pool. These replicates are needed to mitigate the effects of an especially strong or weak training set, which could be selected by chance, especially with the smaller training set sizes. We reported the median metrics of these five replicates.

### Mutational and positional extrapolation

We tested the ability of the models to generalize to mutations and positions not present in the training data, using two dataset splitting strategies referred to as mutational and positional extrapolation. For each of these splitting strategies, we created six replicate train-tune-test splits and tuned the learning rate and batch size independently for each split. We report the Pearson’s correlation on the test set for each split in Figures 2d and S5.

For mutational extrapolation, we designated 80% of single mutations present in the dataset as training and 20% as testing. We then divided the variants into three pools: training, testing, or overlap, depending on whether the variants contained only mutations designated as training, only mutations designated as testing, or mutations from both sets. We discarded the variants in the overlap pool to ensure there was no informational overlap between the training and testing data. We split the training pool into a 90% training set and a 10% tuning set. We used 100% of the variants in the testing pool as the test set.

For positional extrapolation, we followed a similar procedure as mutational extrapolation. We designated 80% of sequence positions as training and 20% as testing. We divided variants into training, testing, and overlap pools, depending on whether the variants contained mutations only in positions designated as training, only in positions designated as testing, or both in positions designated as training and testing. We discarded the variants in the overlap pool, split the training pool into a 90% training set and a 10% tuning set, and used 100% of the variants in the testing pool as the test set.

### Comparison to EVmutation and DeepSequence

We generated multiple sequence alignments using the EVcouplings web server [60] according to the protocol described for EVmutation [1]. We used Jackhmmer [61] to generate an initial alignment with five search iterations against UniRef100 [62] and a sequence inclusion threshold of 0.5 bits/residue. If the alignment had < 80% sequence coverage, we increased the threshold in steps of 0.05 bits/residue until coverage was ≥ 80%. If the number of effective sequences in the alignment was < 10 times the length of the sequence, we decreased the threshold until the number of sequences was ≥ 10 times the length. If the objectives were conflicting, we gave priority to the latter. We set all other parameters to EVcouplings defaults. We trained EVmutation via the “mutate” protocol from EVcouplings. We executed EVmutation locally using EVcouplings v0.0.5 with configuration files generated by the EVcouplings web server. We trained DeepSequence using the same fixed architecture and hyperparameters described in the original work [2]. We fit a DeepSequence model to each alignment and calculated the mutation effect prediction using 2,000 evidence lower bound samples.

### Comparison to Rosetta

We computed Rosetta scores for every variant using Rosetta’s FastRelax protocol with the talaris2014 score function (Rosetta v3.10). First, we created a base structure for each wild-type protein. We generated 10 candidate structures by running relax on the same structure used to generate the protein structure graph, described above. We selected the lowest energy structure to serve as the base. Next, we ran mutate and relax to generate a single structure and compute the corresponding energy for each variant. We set the residue selector to a neighborhood of 10Å. We took the negative of the computed energies to compute the final score for each variant.

### GB1 resampling experiment

We performed a resampling experiment on the GB1 dataset to assess how the quality of deep mutational scanning-derived functional scores impacts performance of supervised learning models. In this case, quality refers to the number of sequencing reads per variant used to estimate the fitness scores. The number of reads per variant depends on the number of variants and the total number of reads in the deep mutational scanning experiment. Raw deep mutational scanning data consists of two sets of variants: an input set (pre-screening) and a selected set (post-screening). Both sets have associated sequencing read counts for each variant, and the functional score for each variant is calculated from these read counts. We resampled the original GB1 data to generate datasets corresponding to 99 different combinations of protein library size and number of reads (Fig. S12). The library size refers to the number of unique variants being screened in the deep mutational scan. Note the final dataset may have fewer unique variants than the protein library. This occurs when there is a low number of sequencing reads relative to the size of the library. In that scenario, not all generated variants will get sequenced, even though they were screened as part of the function assay.

First, we created a filtered dataset by removing any variants with zero reads in either the input set or selected set of the original deep mutational scanning data. We generated Enrich2 scores for this filtered dataset using the same approach described in the above section on datasets. We randomly selected 10,000 variants from this dataset to serve as a global testing set. Next, we used the filtered dataset, minus the testing set variants, as a base to create the resampled datasets. For each library size in the Figure 3 heat map, we randomly selected that many variants from the base dataset to serve as the library. Then for each library, we created multinomial probability distributions giving the probability of generating a read for a given variant. We created these probability distributions for both the input and selected sets by dividing the original read counts of each variant by the total number of reads in the set. The multinomial distributions allowed us to sample new input and selected sets based on the read counts in the Figure 3 heat map. To determine how many reads should be sampled from the input set versus the selected set, we computed the fraction of reads in the input set and selected set in the base dataset and sampled reads based on that fraction. Finally, we generated Enrich2 scores for each resampled dataset using the same approach described in the above section on datasets. To account for potential skewing from random sampling, we generated five replicates for each of the 99 combinations of library size and numbers of reads. Counting the replicates, we created 495 resampled datasets in total.

We trained the supervised learning models on each resampled dataset, as long as the dataset had at least 25 total variants in each of its 5 replicates. Out of the 99 combinations of library size and number of reads, 7 did not have enough variants across the replicate datasets and were thus excluded from this experiment. Although the libraries of these 7 combinations had more than 25 variants, there were not enough reads to estimate scores for all of them, and thus the final datasets ended up with less than 25 variants. We split each resampled dataset into 80% training and 20% tuning sets. The tuning sets were used to select the learning rate and batch size hyperparameters. The network architectures and other parameters were set to those selected during the main experiment described above. We evaluated each model using the held-out testing set with non-resampled fitness scores. This type of evaluation ensures that although the models are trained on resampled datasets with potentially unreliable fitness scores, they are evaluated on high-confidence fitness scores from the non-resampled dataset. We report the mean Pearson’s correlation coefficient across the five replicates for each combination of library size and number of reads.

### UMAP projection of latent space

Each neural network encodes a latent representation of the input in its last internal layer before the output node. The last internal layer in the convolutional networks is a dense fully connected layer with 100 hidden nodes. Thus, the latent representation of each variant at this layer is a length 100 vector. We used UMAP [22] to project the latent representation of each variant into a two-dimensional space to make it easier to visualize while still preserving spatial relationships between variants. We used the umap-learn package v0.4.0 to compute the projection with default hyperparameters (n_neighbors=15, min_dist=0.1, and metric=“euclidean”). The two-dimensional visualization shows how the network organizes variants internally prior to predicting a functional score. We colored each variant by its score to show that the network efficiently organizes the variants. Variants grouped close together in the UMAP plot have similar functional scores. We also annotated a few key variants, such as the highest and lowest scoring variants.

### Integrated gradients

To determine which input features were important for making predictions, we generated integrated gradients feature attributions [23] for all variants. The attributions quantify the effects of specific feature values on the network’s output. A positive attribution means the feature value pushes the network to output a higher score relative to the given baseline, and a negative attribution means the feature value pushes the network to output a lower score relative to the given baseline. We used the wild-type sequence as the baseline input. Integrated gradients attributions are computed on a per-variant basis, meaning attributions are specific to the feature values of the given variant. Due to nonlinear effects captured by the nonlinear models, a given feature value might have a positive attribution in one variant but a negative attribution in a different variant. We computed attributions for all variants in the training set. Examining the training set is analogous to other model interpretation techniques that compute attributions directly from the weights or parameters of models that were trained using training sets. We summed the attributions for all features at each sequence position, allowing us to see which mutations pushed the network to output a higher or lower score for each individual variant. We also summed the attributions across all the variants in the training set to see which sequence positions were typically tolerant or intolerant to mutations. We used DeepExplain [63] v0.3 to compute the integrated gradients attributions with “steps” set to 100.

### Model-guided design of GB1 variants

We used a random-restart hill climbing algorithm to design sequences with a set number of mutations (*n*) from wild-type GB1 that maximized the minimum predicted functional score from an ensemble of four models (linear regression, fully connected, sequence convolutional, and graph convolutional).

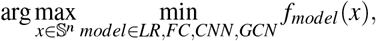

where *x* is a sequence, 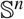 is the set of all sequences *n* mutations from wild-type, and *f_model_*(*x*) is a model’s predicted score for sequence *x*. This design objective ensures that all models predict the sequence will have a high functional score. We initialized a hill climbing run with a randomly selected sequence containing *n* point mutations and performed a local search by exchanging each of these *n* mutations with each other possible single point mutation. Exchanging mutations ensured that we only search sequences a fixed distance *n* from wild type. We then moved to the mutation-exchanged variant with the highest objective, which became our new reference point, and repeated this hill climbing process until a local optima was reached. We performed each sequence optimization with ten random initializations and took the design with the highest overall objective value. We applied this procedure to design one sequence at each level of diversity, where *n* = 10, 20, 30, 40, 50. We visualised the sequence space using multidimensional scaling with the Hamming distance as a distance metric between sequences.

We predicted the three-dimensional structure of Design10 using Rosetta Abinitio [64]. We used the Rosetta Fragment Server to generate the fragments for the Design10 sequence. We generated 100 structures using Rosetta 3.12 and AbinitioRelax and selected the structure with the lowest total score. The predicted structure for Design10 aligns to the wild-type GB1 crystal structure with 0.9 Å C_*α*_ root-mean-square deviation.

### Designed GB1 variant gene synthesis and protein expression

We designed the genes encoding the designed GB1 variants by making codon substitutions into the base wild-type GB1 gene sequence. If there were multiple codon options for an amino acid, we chose the particular codon randomly from a set of 31 codons that are optimized for expression in E. coli [65]. For our expression construct, we included an upstream bicistronic design (BCD) element to minimize any influence of mRNA secondary structure on protein expression [66] and also included an N-terminal 6x His-tag with a five-amino-acid linker for protein purification. We ordered wild-type GB1 and the five designed GB1 variants from Twist Biosciences cloned into the pET21(+) protein expression vector.

We expressed wild-type GB1 and the five designed GB1 variants using a standard T7 expression system. We transformed the six plasmids into BL21(DE3) E. coli cells. We expressed the GB1 variants by inoculating LB cultures containing 100 *μ*g/mL carbenicillin with a 1:100 dilution of overnight cultures, incubating these cultures shaking at 37°C until they reached an OD600 of 0.4-0.6, and inducing with 400 *μ*M Isopropyl *β*-D-1-thiogalactopyranoside (IPTG). We then incubated these expression cultures overnight at 20°C while shaking, pelleted the cells by centrifuging at 3000 g for 20 minutes at 4°C, and stored the cell pellets at −80°C.

We determined the level of soluble protein expression using sodium dodecyl sulphate–polyacrylamide gel electrophoresis (SDS-PAGE). We thawed the cell pellets on ice and resuspended into 0.5 mL of Buffer A (20 mM sodium phosphate pH 7.3, 500 mM NaCl, 20 mM imidazole). We then added 2.5 mL of lysis buffer (Buffer A + 0.60x BugBuster + 2 U/mL DNaseI (Thermo Fischer) + 1 mg/mL hen egg white lysozyme) to each sample and incubated at room temperatures for 5 minutes to yield the total cell lysate. We obtained the soluble protein fraction by centrifuging at 21,000 g for 70 minutes and extracting the supernatant. We then ran samples of the total cell lysate and soluble fractions on a Novex 4-20% Tris-Glycine SDS-PAGE gel (Thermo Fischer). After staining, we analyzed the gels to qualitatively evaluate whether the expressed proteins were present in the soluble fraction

### Protein purification and circular dichroism spectroscopy

We expressed wild-type GB1 and Design10 using the above protocol, with the exception that the expression cultures were incubated at 16°C for 24 hours. We thawed the cell pellets on ice, resuspended in 2.5 mL of Buffer A, sonicated for 1 minute with 5 second pulses spaced by 15 second resting periods, and centrifuged for 10 minutes at 21,000 g to obtain the soluble protein fraction. We then ran the soluble fraction over a Ni Sepharose 6 Fast Flow column (Cytiva Life Sciences) that was equilibrated with Buffer A, washed with 3 column volumes of Buffer A, and eluted in 1.5 mL fractions of Buffer B (20 mM sodium phosphate pH 7.3, 500 mM NaCl, 500 mM imidazole). We ran the elution fractions over SDS-PAGE and pooled the fractions that contained the target protein. Finally, we aliquoted the purified protein, flash froze the aliquots in liquid nitrogen, and stored at −80°C.

For circular dichoism (CD) spectroscopy, we thawed the purified protein samples on ice and dialyzed overnight in 20 mM sodium phosphate pH 8.0 at 4°C to remove imidazole. We then determined the protein concentrations using a Nanodrop spectrophotometer. The CD measurements were then performed by UW-Madison’s Biophysics Instrumentation Facility. They measured CD spectra using a 1 mm pathlength on an AVIV Model 420 Circular Dichroism Spectrometer at 4°C. The CD spectra for Design10 was normalized to the wild-type GB1 spectra at 222 nm.

### GB1 yeast display plasmid construction and flow cytometric IgG binding affinity titration

We synthesized wild-type and Design10 variant GB1 genes as yeast codon-optimized gBlocks (Integrated DNA Technologies, Coralville, IA). The gBlocks were ligated into the unique NheI and BamHI sites of the yeast surface display vector pCTCON2 (provided by Dane Wittrup, MIT). We synthesized A24Y and E19Q+A24Y variant GB1 genes as yeast-optimized gene fragments (Twist, San Francisco, CA). The gene fragments were ligated into a golden-gate compatible version of pCTCON2 at the NheI, BamHI sites. This yeast display vector fuses the Aga2p protein to the N-terminus of GB1.

We transformed plasmid DNA into yeast display *Saccharomyces cerevisiae* strain EBY100 made competent using the Zymo Research Frozen EZ Yeast Transformation II kit with transformants grown on synthetic dropout (SD) -Trp (MP Biomedicals, Irvine, CA) agar plates for two days at 30°C. After two days, individual colonies were picked into 4 mL of low-pH Sabouraud Dextrose Casamino Acid media (20 g/L dextrose, 6.7 g/L yeast nitrogen base, 5 g/L casamino acids, 10.4 g/L sodium citrate, 7.4 g/L citric acid monohydrate) and grown overnight at 30°C and 250 rpm. For induction of GB1 display, we started a 5 mL Sabouraud Galactose Casamino Acid (8.6 g/L Na_2_PO*H_2_O, 5.4 g/L Na_2_HPO_4_, 20 g/L galactose, 6.7 g/L yeast nitrogen base, 5 g/L casamino acids) culture at an optical density, as measured at 600 nm, of 0.5 and shook overnight at 250 rpm and 20°C.

We harvested approximately 2 × 10^5^ yeast cells for each titration data point by centrifugation after overnight incubation, washed them once in pH 7.4 Phosphate Buffered Saline (PBS) containing 0.2% (w/v) bovine serum albumin (BSA), and incubated them overnight at 4°C on a tube rotator at 18 rpm in between 100 *μ*L and 800 *μ*L of PBS/0.2% BSA containing various concentrations of mouse IgG2a (BioLegend, San Diego, CA) that had been conjugated with Alexa647 using NHS chemistry (Molecular Probes, Eugene, OR). Volumes of Alexa647 IgG-containing incubation solution were varied to prevent ligand depletion from occurring in the lowest IgG concentration incubation tubes. Following overnight incubation, yeast were washed once in PBS/0.2% BSA and resuspended in ice cold PBS for flow cytometric analysis. Analyses were performed using a Fortessa analyzer (Becton Dickinson), and the mean of the fluorescence distribution was reported.

We performed duplicate fluorescence measurements for all nine IgG concentrations tested. We then fit a Hill function to the average of these duplicate measurements. We were able to determine the *K_d_* of Design10 as 5 nM because the binding curve was beginning to display saturation. We were unable to determine the *K_d_* of wild-type, A24Y, or E19Q+A24Y GB1 variants because the proteins were less than 50% bound at the highest IgG concentration tested.

### Code availability

We provide a cleaned version of our code that can be used to retrain the models from this publication or train new models with different network architectures or for different datasets. We also provide pre-trained models that use the latest code and are functionally equivalent to the ones from this publication. The pre-trained models can be used to make predictions for new variants. Our code is freely available on GitHub and is licensed under the MIT license: https://github.com/gitter-lab/nn4dms. The software is also archived on Zenodo at https://doi.org/10.5281/zenodo.4118330. See Table S5 for software dependencies and their versions.

## Acknowledgements

We thank Zhiyuan Duan for his assistance running Rosetta and Darrell McCaslin at the University of Wisconsin-Madison Biophysics Instrumentation Facility for his expertise in collecting and analyzing the circular dichroism spectra.

## Supplementary Information

**Figure S1.**
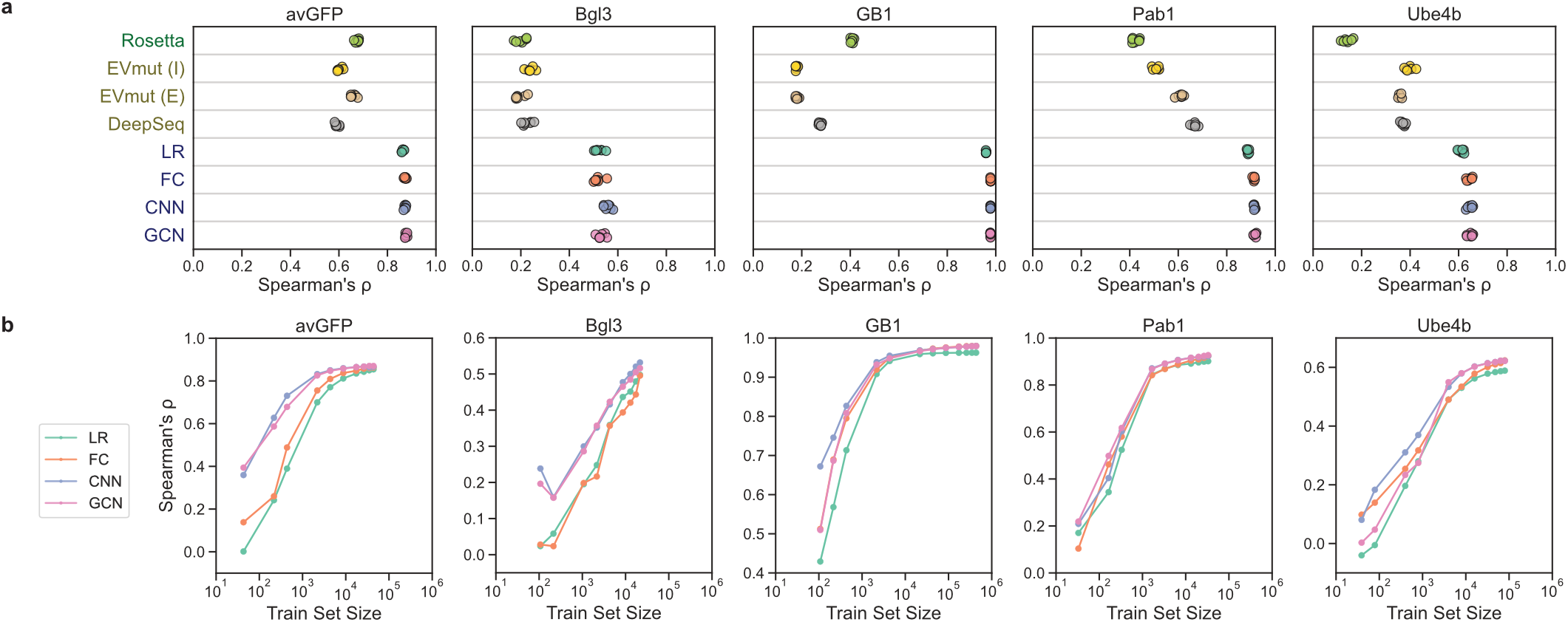
Model evaluation using Spearman’s correlation coefficient. (a) Spearman’s correlation coefficient between true and predicted scores for Rosetta, EVmutation, DeepSequence, linear regression (LR), fully connected network (FC), sequence convolutional network (CNN), and graph convolutional network (GCN). EVmutation (I) refers to the independent formulation of the model that does not include pairwise interactions. EVmutation (E) refers to the epistatic formulation of the model that does include pairwise interactions. Each point corresponds to one of seven random train-tune-test splits. (b) Spearman’s correlation performance of supervised models trained with reduced training set sizes.

**Figure S2.**
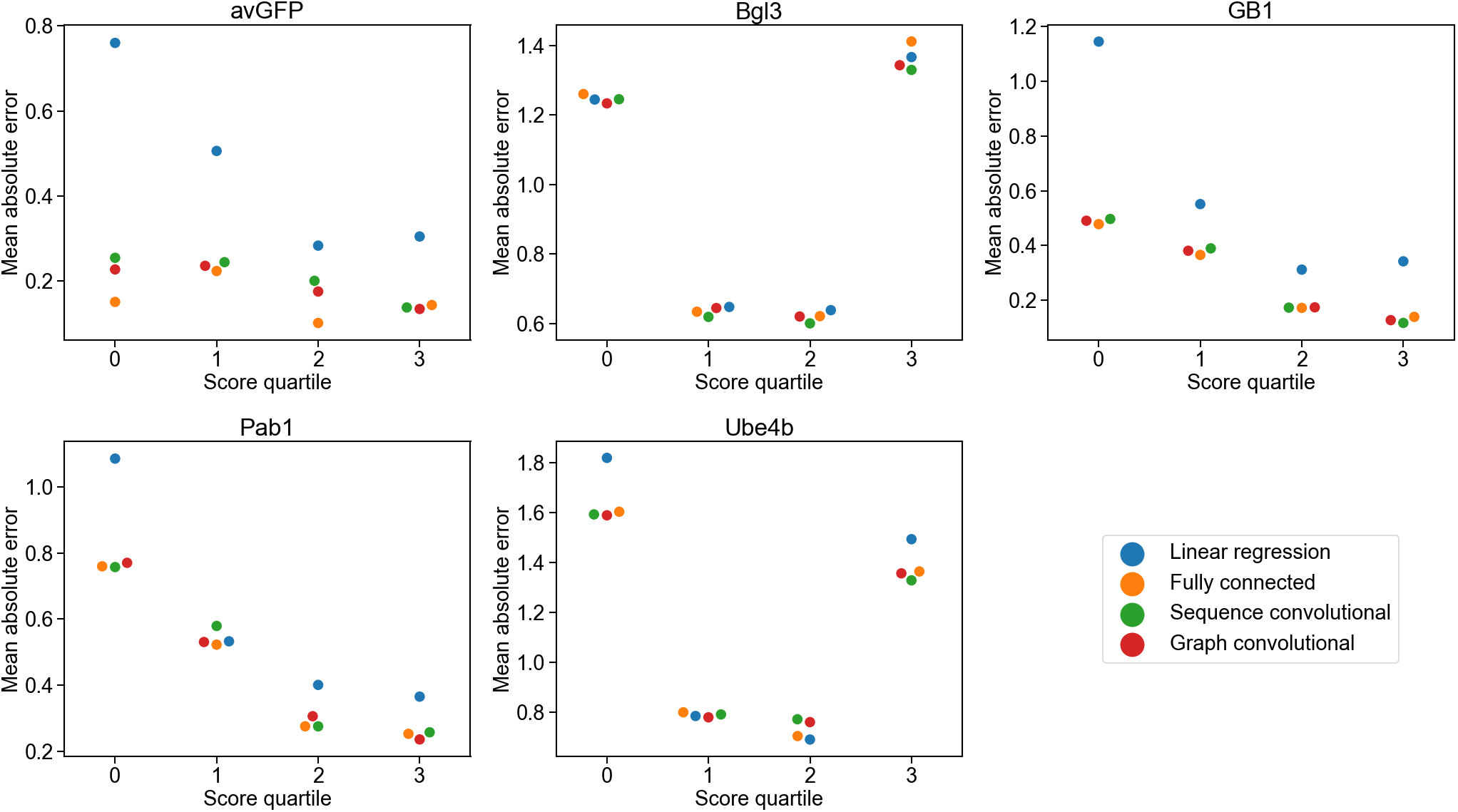
Mean absolute error vs. score quartile. The mean absolute error in the models’ predictions grouped by score quartile. Linear regression has a substantial jump in error for low-scoring variants compared to the other models in avGFP, GB1, Pab1, and Ube4b.

**Figure S3.**
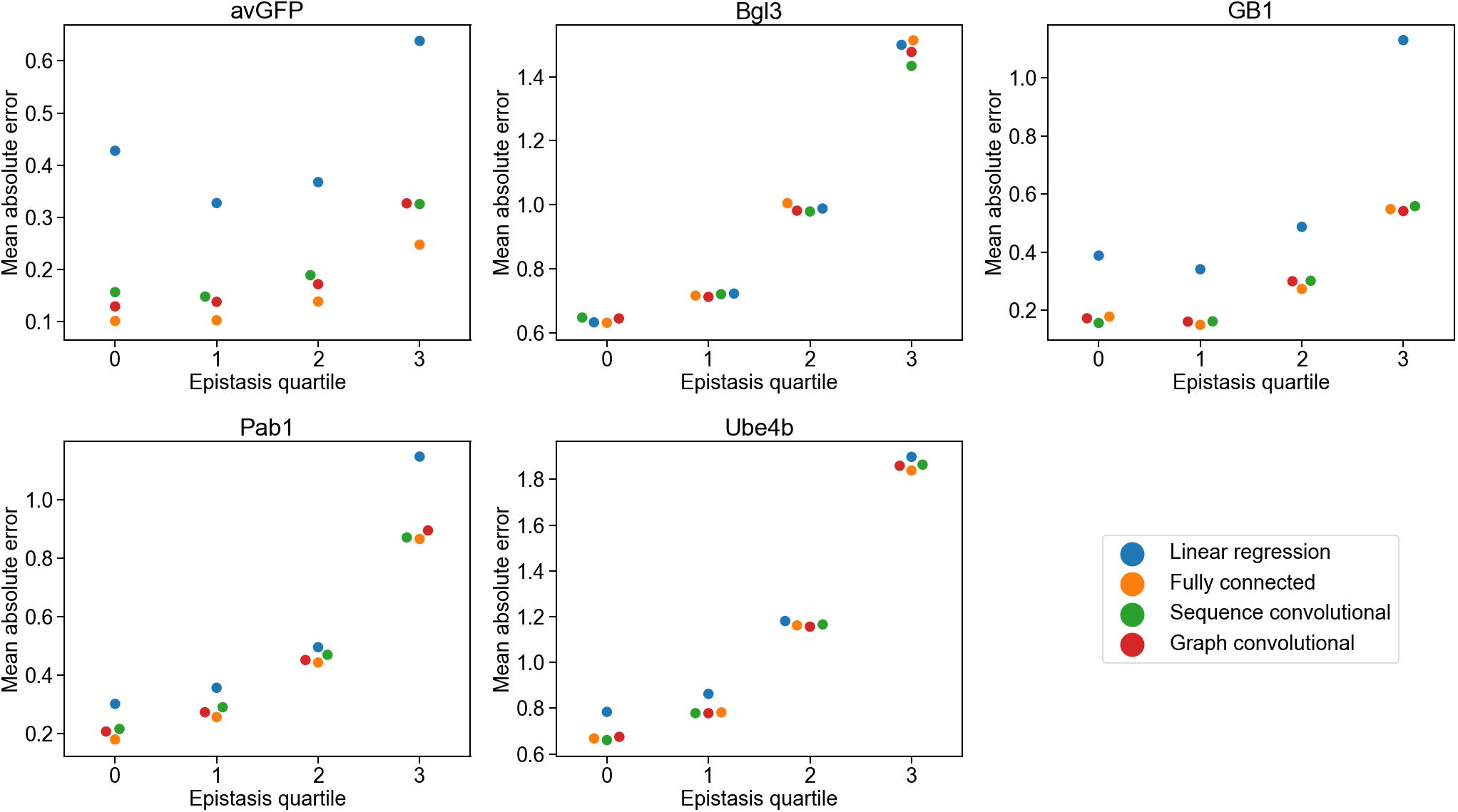
Mean absolute error vs. epistasis quartile. The mean absolute error in the models’ predictions grouped by absolute epistasis quartile. We compute epistasis by subtracting the expected score for the multi-mutant sequence from the true score. The expected score for the multi-mutant sequence is the sum of the corresponding single-mutant scores, truncated to the observed minimum or maximum in the dataset. Linear regression has a substantial jump in error for high-epistasis variants compared to the other models in avGFP, GB1, and Pab1.

**Figure S4.**
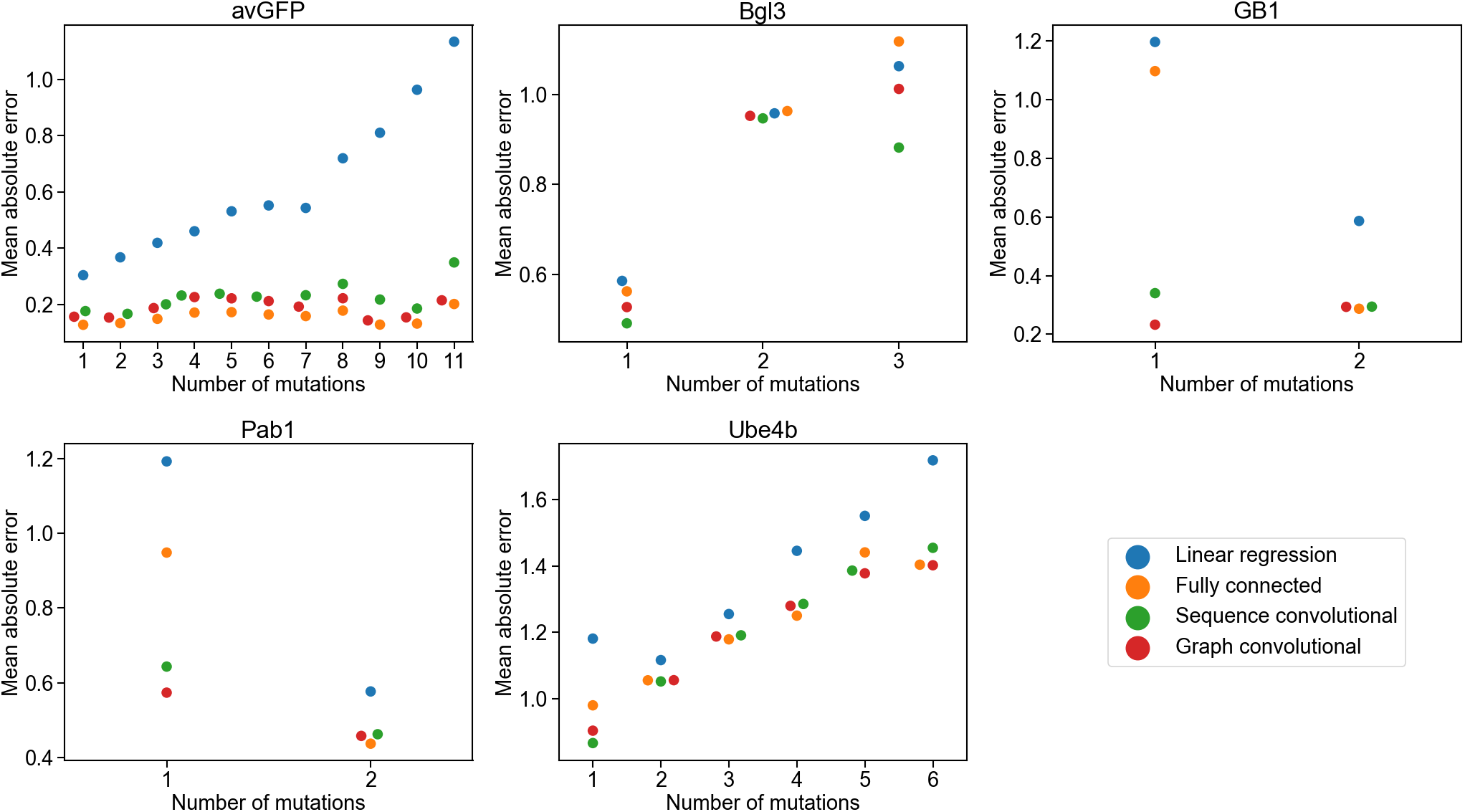
Mean absolute error vs. number of mutations. The absolute error in the models’ predictions for each variant grouped by the number of mutations in the variant. Linear regression struggles with increasing numbers of mutations in avGFP. The convolutional networks perform better than linear regression and the fully connected network on single-mutation variants in GB1 and Pab1.

**Figure S5.**
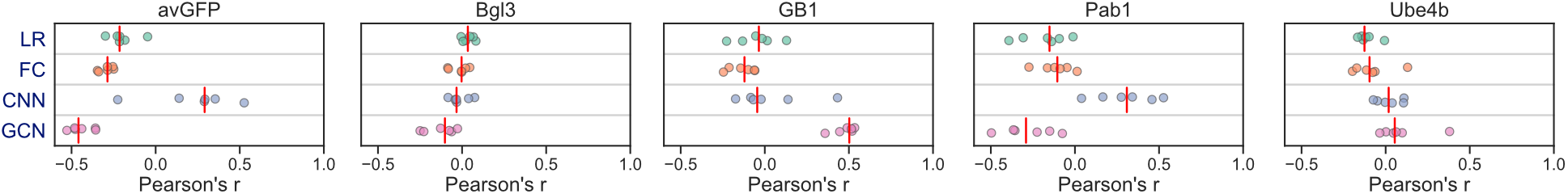
Positional extrapolation. Model performance when making predictions for variants containing mutations in positions that were unmodified in the training data (positional extrapolation). Each point corresponds to one of six replicates, and the red vertical line denotes the median.

**Figure S6.**
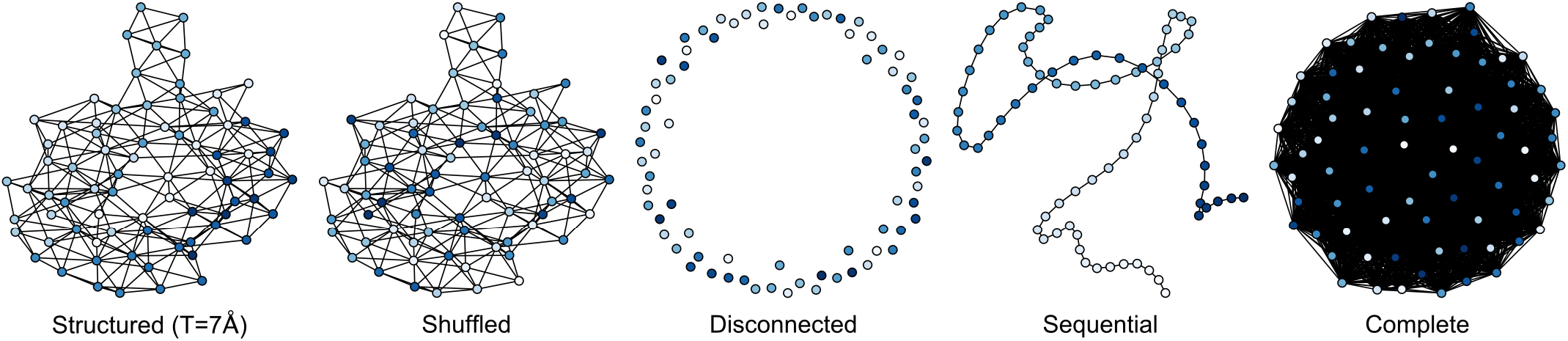
Protein structure graphs for Pab1. The graph convolutional network uses a graph of the protein’s structure to determine which residues are close together. In addition to the standard graph based on the protein’s actual structure, we tested four baseline graphs: a shuffled graph based on the standard graph but with shuffled node labels, a disconnected graph with no edges, a sequential graph containing only edges between sequential residues, and a complete graph containing all possible edges. The graphs pictured are for the Pab1 dataset. The structured graph uses a distance threshold of 7Å to determine which residues should be connected with edges (selected by hyperparameter sweep). The nodes are colored according to each residue’s sequence position, with light colors corresponding to residues at the start of the sequence and dark blue colors corresponding to residues at the end of the sequence.

**Figure S7.**
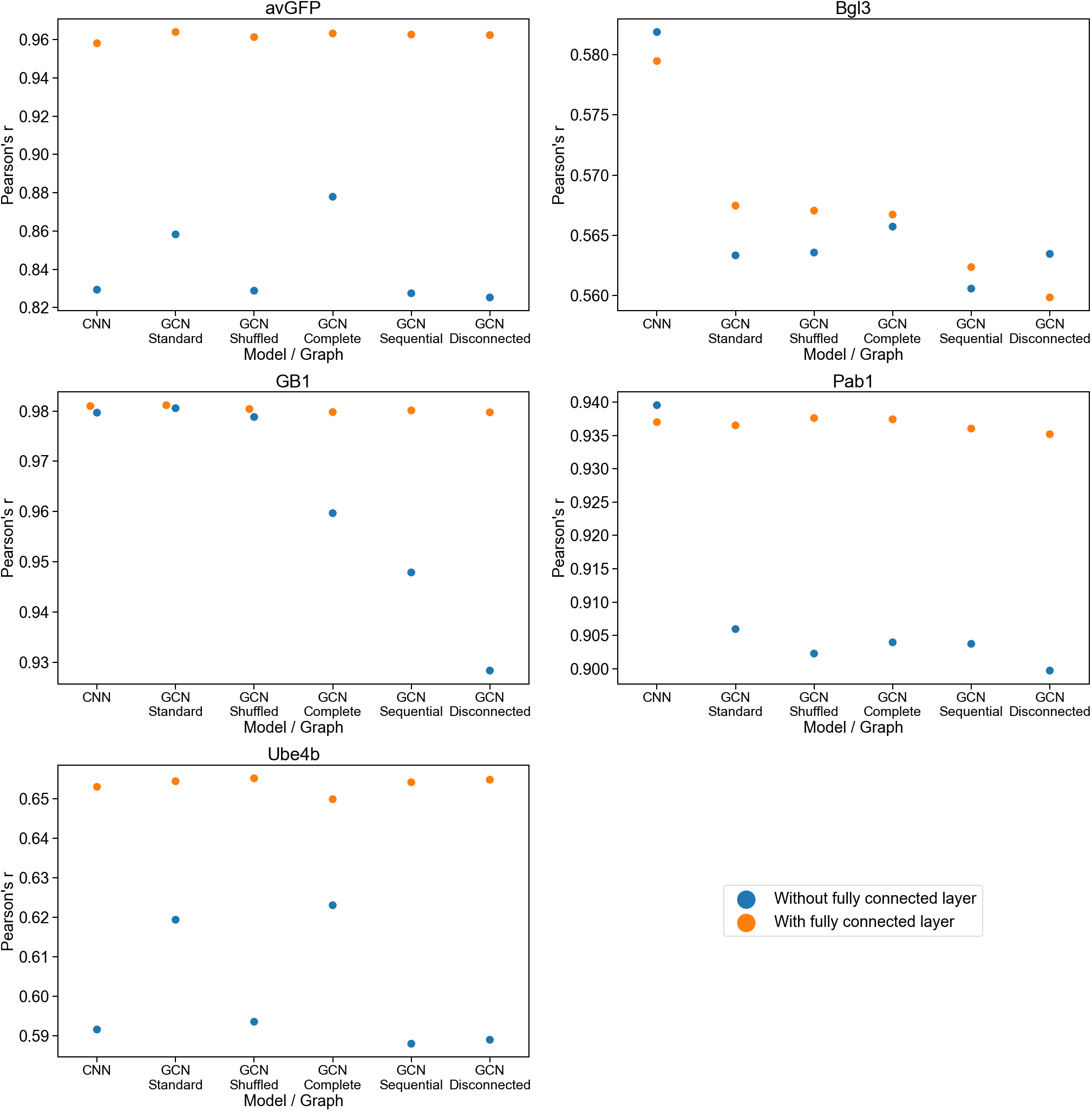
Convolutional networks with and without a fully connected layer. The correlation performance of sequence convolutional and graph convolutional networks trained with various baseline structure graphs, with and without a final fully connected layer. The standard graph is based on the protein’s actual structure. The shuffled graph is a version of the regular structured graph with shuffled node labels. The complete graph contains all possible edges between residues. The sequential graph only contains edges between sequential residues. The disconnected graph contains no edges. The fully connected layer at the end of the network compensates for apparent differences in performance caused by type of convolutional network or different graph structures.

**Figure S8.**
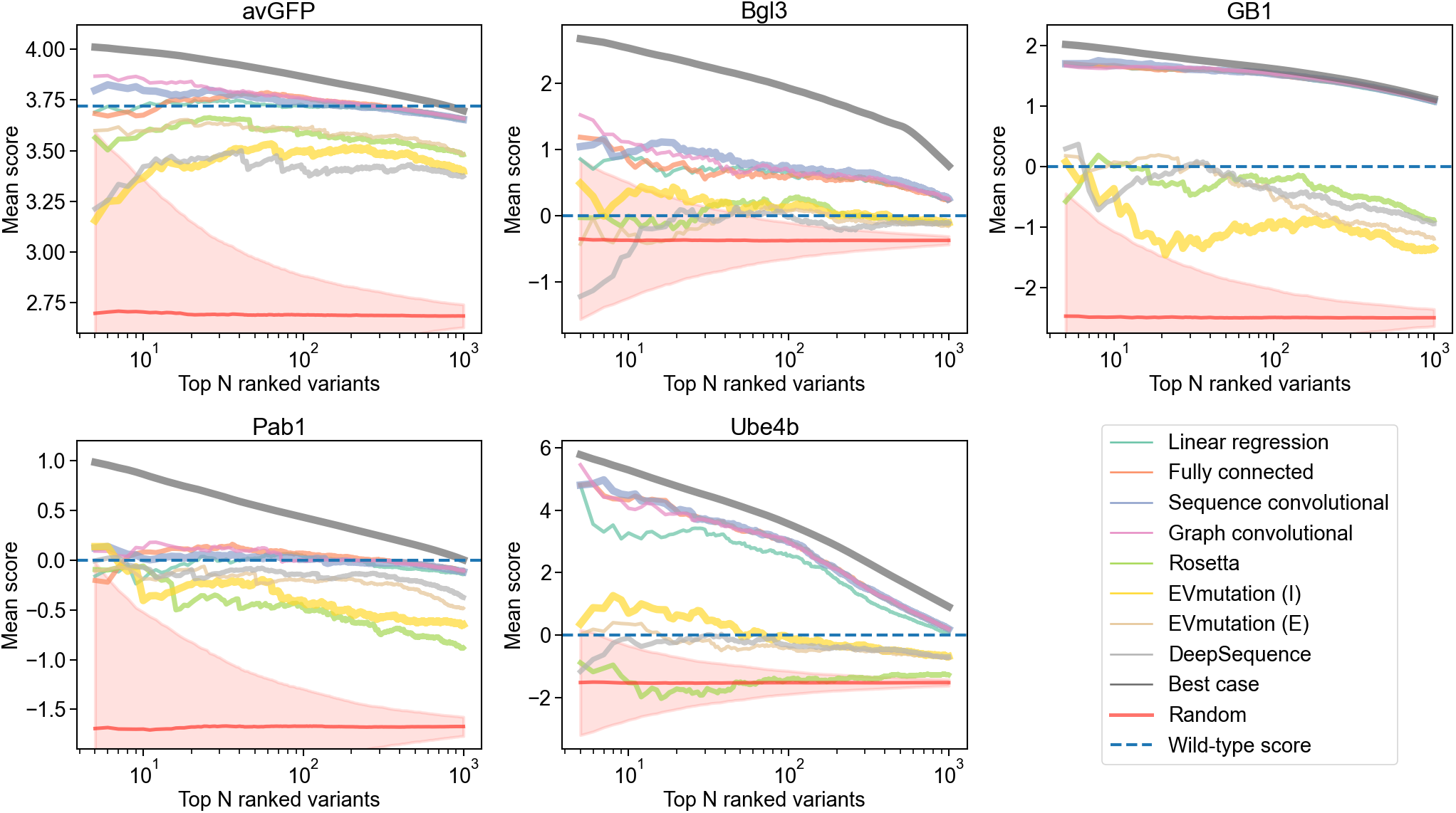
Mean score of highest ranked variants. The mean score of each model’s ranking of the highest scoring test set variants. For the most part, the supervised models prioritize variants whose average score is higher than the wild-type. The random baseline is shown with the mean and 95% confidence interval.

**Figure S9.**
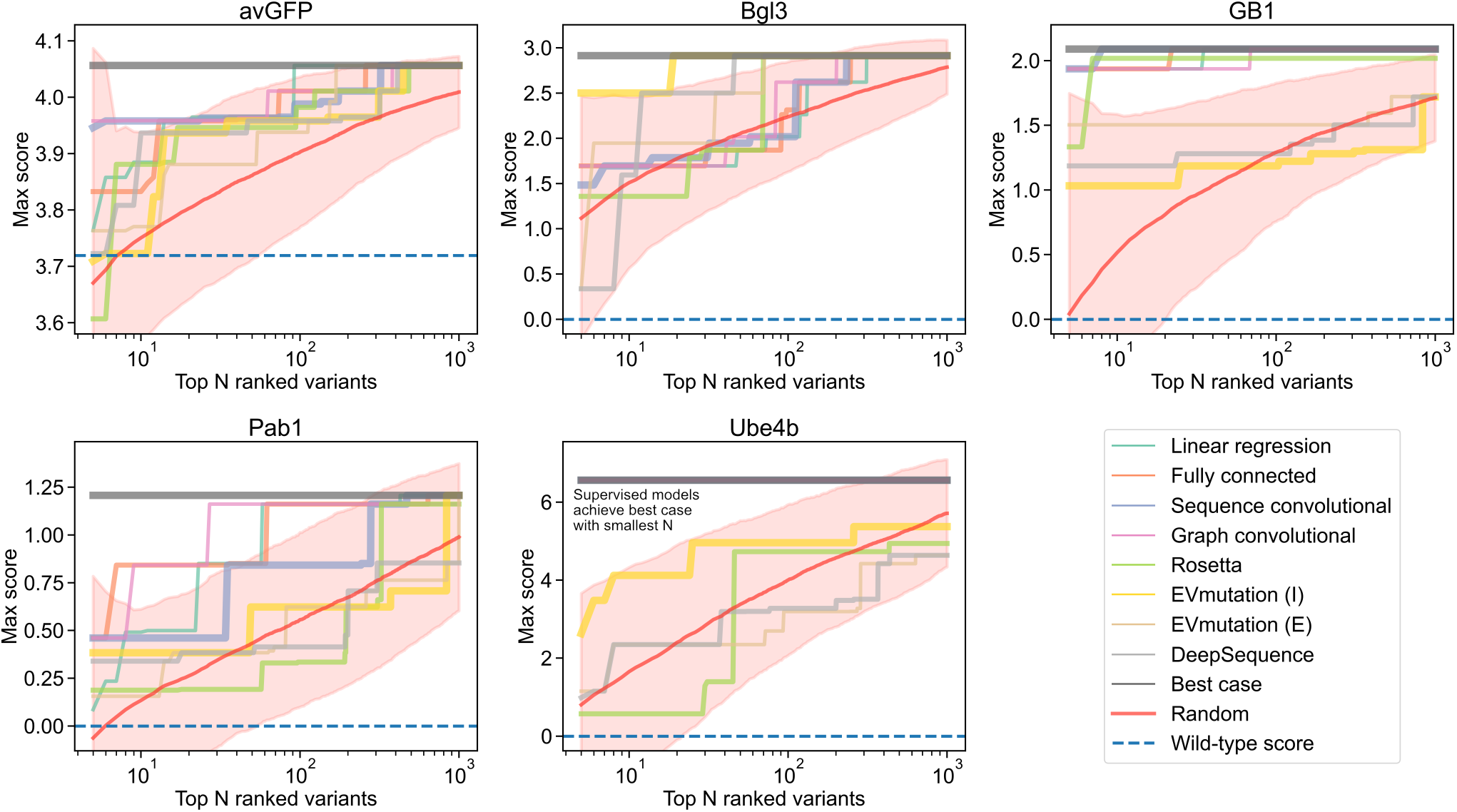
Max score of highest ranked variants. The max score in each model’s ranking of the highest scoring test set variants. For Ube4b, the supervised models prioritize a variant with the true max score with the smallest tested budget (N=5), thus all the lines corresponding to the supervised models are hidden behind the line for the true score. Nearly all models across all datasets prioritize variants whose max score is higher than the wild-type. The random baseline is shown with the mean and 95% confidence interval.

**Figure S10.**
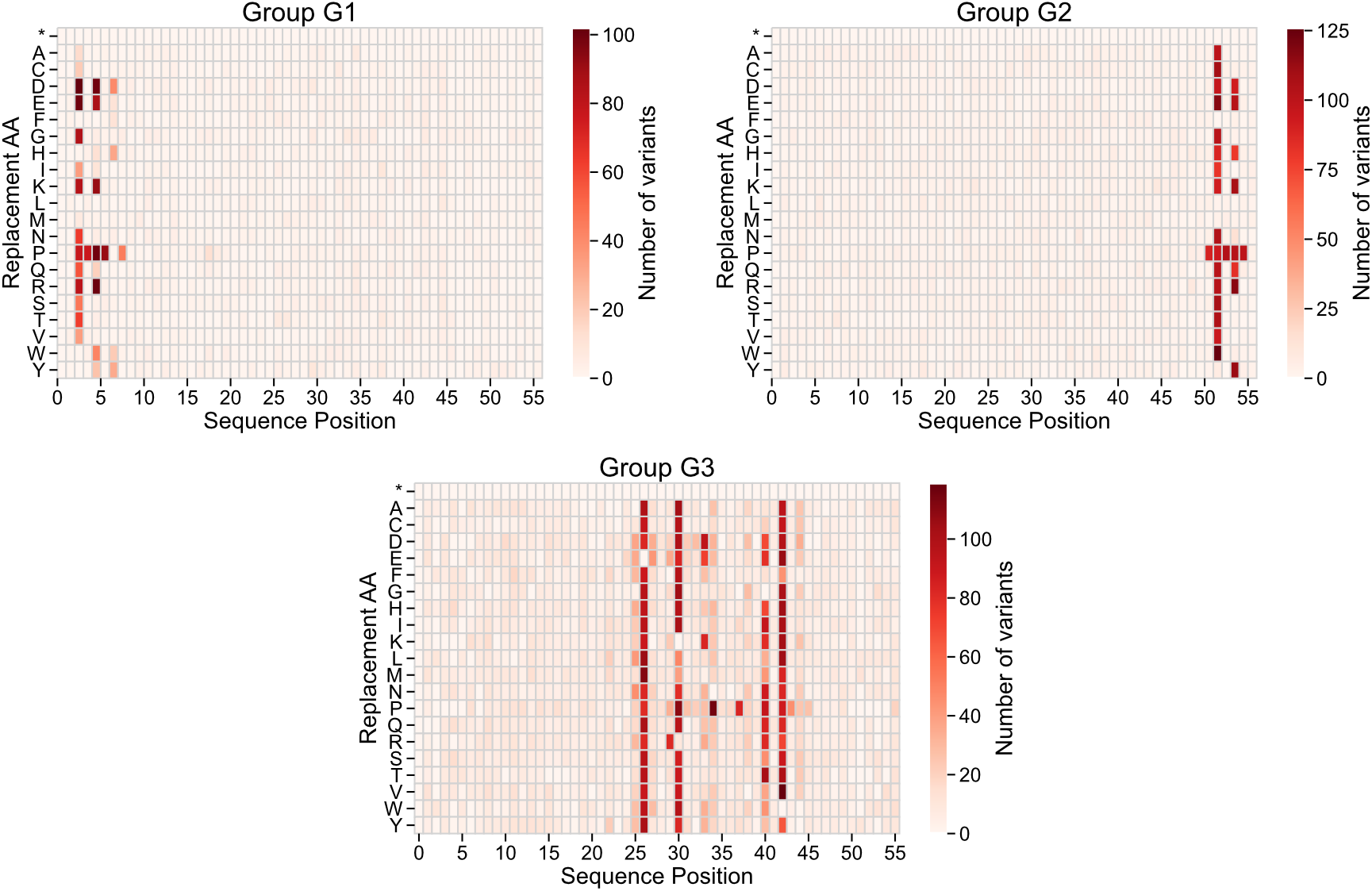
Mutations in GB1 latent space groups. Heat maps showing the number of occurrences of mutations for each annotated group in the GB1 latent space in Figure 4a. Groups G1 and G2 contain variants with mutations at core residues near the start and end of the sequence, respectively. Group G3 contains variants with mutations at surface interface residues.

**Figure S11.**
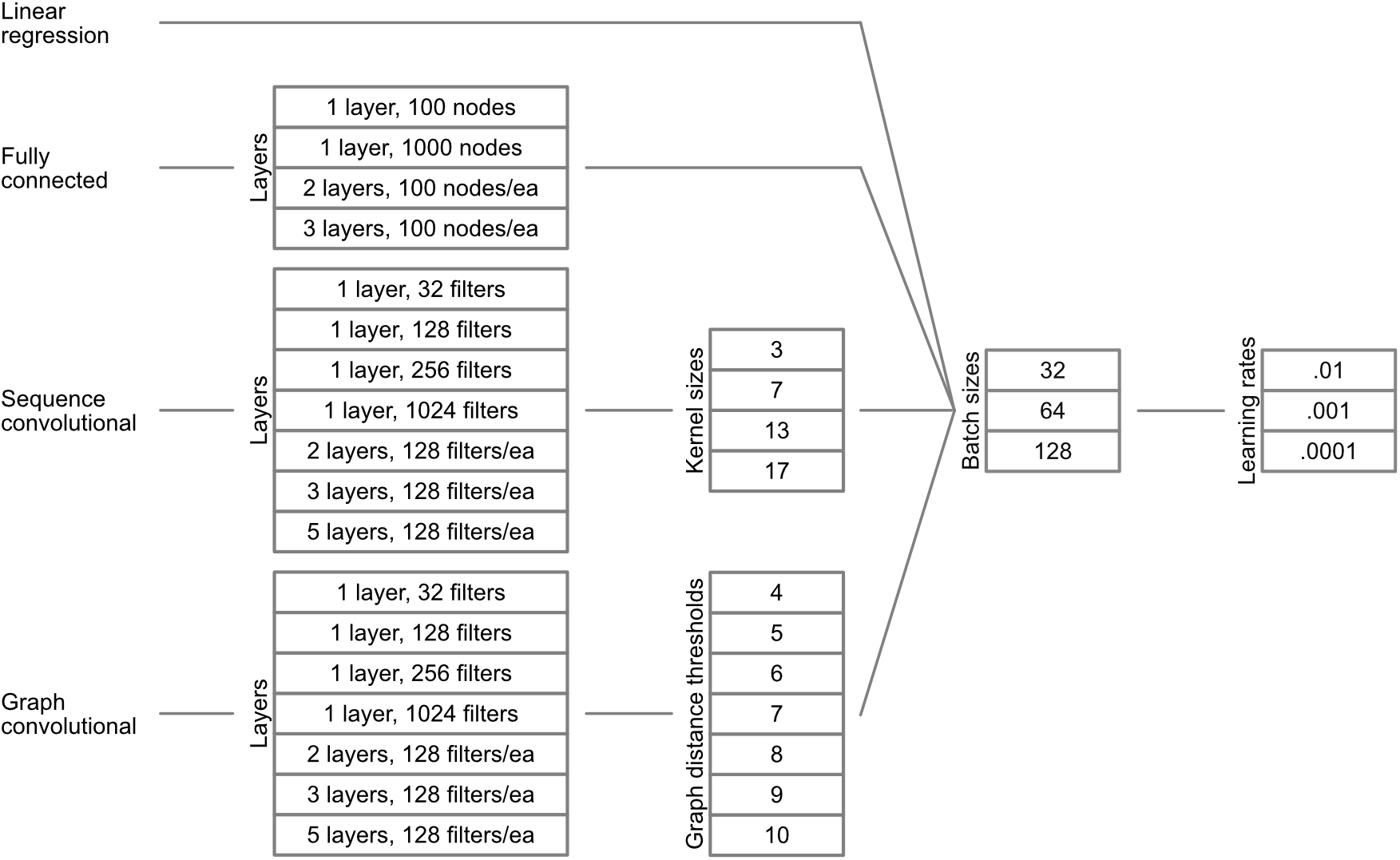
Hyperparameter sweep. We performed an exhaustive hyperparameter sweep for each dataset and type of model using all possible combinations of these hyperparameters.

**Figure S12.**
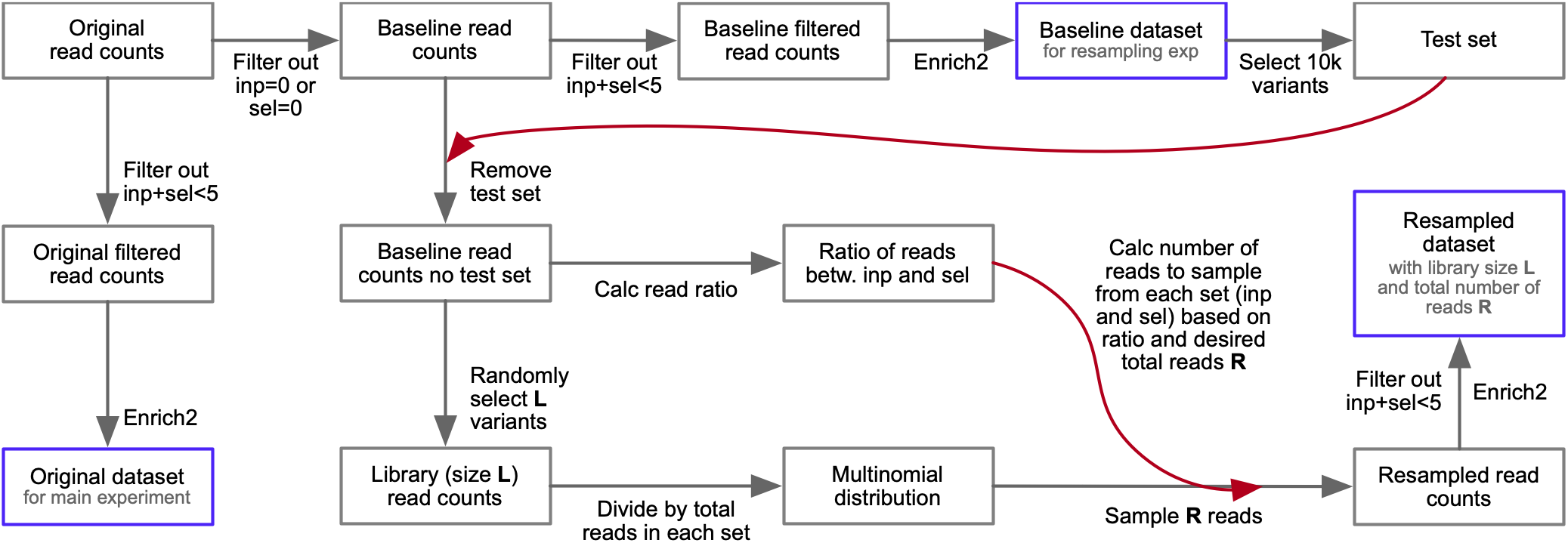
Generation of resampled GB1 datasets. Flowchart showing how we created resampled GB1 datasets corresponding to different library sizes and numbers of reads.

**Table S1.**
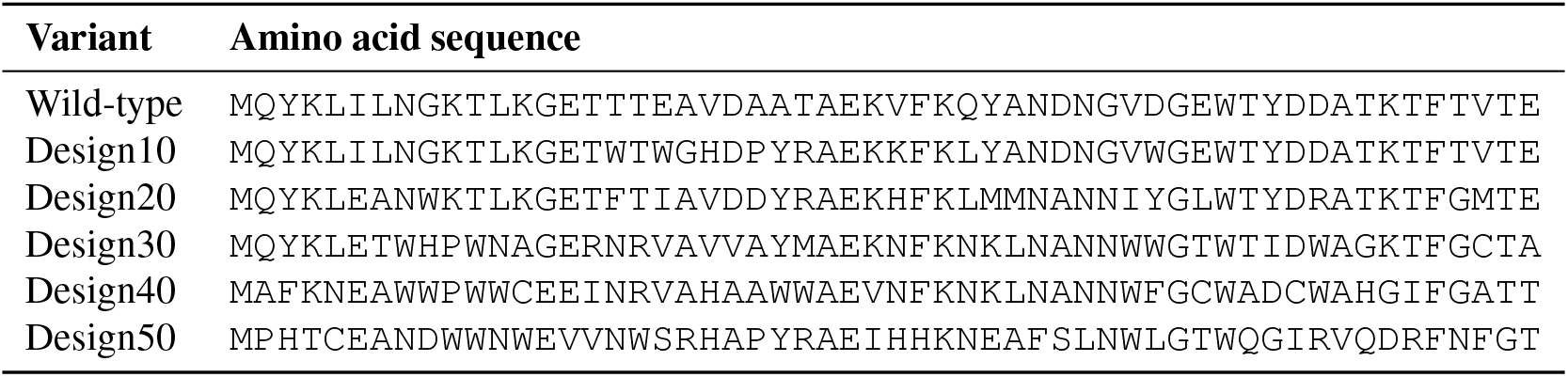
Designed GB1 sequences. The GB1 wild-type sequence and the designed sequences with increasing numbers of mutations (10, 20, 30, 40, and 50) from wild-type.

**Table S2.** Diversity in designed GB1 sequences. We repeated our hill climbing protein design approach 100 times to generate 100 sequences with 10 mutations each. We found 27 of 100 design runs converged to the same sequence. The other 73 represent distinct local optima in the landscape. A number of mutations were observed across multiple designs, and some of these were present in Design10. This table lists mutations common across the designs and their frequencies.

| Mutation | Frequency | Present in Design10 |
| --- | --- | --- |
| A24Y | 0.93 | Y |
| D40W | 0.52 | Y |
| D40Y | 0.48 | N |
| V29K | 0.48 | Y |
| Q32L | 0.45 | Y |
| E42Q | 0.42 | N |
| A34M | 0.33 | N |

**Table S3.** Selected hyperparameters. The hyperparameters selected by a hyperparameter sweep for the main experiment. There are additional parts of the architecture that were not part of the hyperparameter sweep. For example, the fully connected networks have a dropout layer after every dense layer. The convolutional networks have a dense layer and a dropout layer before the output node. Experiments with reduced training set sizes and GB1 resampling used the same architectures selected for the main experiment, but they had their own sweeps for learning rate and batch size.

| Dataset | Model type | Key parts of architecture | Learning rate | Batch size | Epochs |
| --- | --- | --- | --- | --- | --- |
| avGFP | Linear regression | Linear regression | 0.0001 | 128 | 90 |
|  | Fully connected | 3 layers, 100 hidden units each | 0.0001 | 32 | 134 |
|  | Sequence convolutional | 5 layers, kernel size 3, 128 filters | 0.001 | 64 | 113 |
|  | Graph convolutional | 2 layers, 7Å threshold, 128 filters | 0.0001 | 32 | 130 |
| Bgl3 | Linear regression | Linear regression | 0.0001 | 128 | 164 |
|  | Fully connected | 2 layers, 100 hidden units each | 0.0001 | 32 | 187 |
|  | Sequence convolutional | 1 layer, kernel size 17, 32 filters | 0.0001 | 64 | 102 |
|  | Graph convolutional | 1 layer, 6Å threshold, 32 filters | 0.0001 | 128 | 129 |
| GB1 | Linear regression | Linear regression | 0.0001 | 128 | 27 |
|  | Fully connected | 1 layer, 1000 hidden units | 0.0001 | 64 | 110 |
|  | Sequence convolutional | 3 layers, kernel size 17, 128 filters | 0.0001 | 32 | 27 |
|  | Graph convolutional | 5 layers, 7Å threshold, 128 filters | 0.0001 | 32 | 109 |
| Pab1 | Linear regression | Linear regression | 0.001 | 128 | 47 |
|  | Fully connected | 3 layers, 100 hidden units each | 0.001 | 128 | 108 |
|  | Sequence convolutional | 3 layers, kernel size 17, 128 filters | 0.0001 | 128 | 42 |
|  | Graph convolutional | 1 layer, 7Å threshold, 32 filters | 0.0001 | 128 | 232 |
| Ube4b | Linear regression | Linear regression | 0.0001 | 64 | 73 |
|  | Fully connected | 3 layers, 100 hidden units each | 0.0001 | 64 | 124 |
|  | Sequence convolutional | 5 layers, kernel size 3, 128 filters | 0.0001 | 128 | 29 |
|  | Graph convolutional | 3 layers, 7Å threshold, 128 filters | 0.0001 | 128 | 92 |

**Table S4.**
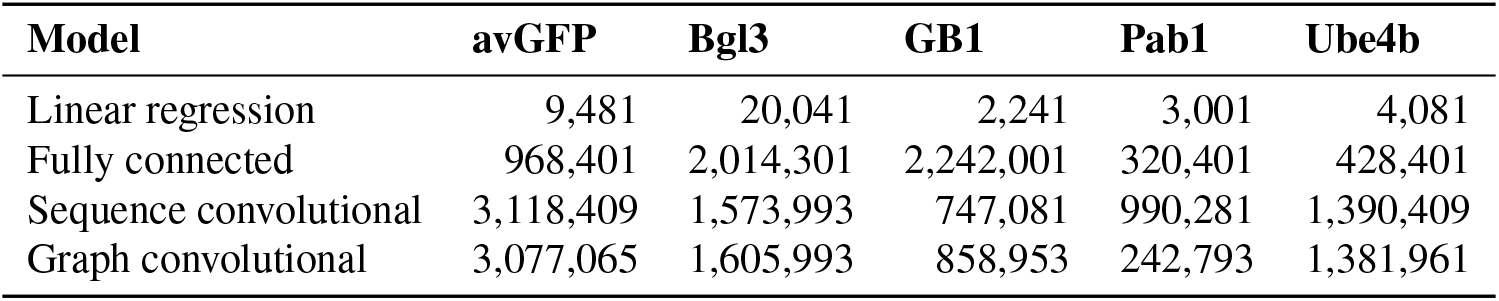
Numbers of trainable parameters. The number of trainable parameters in each model.

| Model | avGFP | Bgl3 | GB1 | Pab1 | Ube4b |
| --- | --- | --- | --- | --- | --- |
| Linear regression | 9,481 | 20,041 | 2,241 | 3,001 | 4,081 |
| Fully connected | 968,401 | 2,014,301 | 2,242,001 | 320,401 | 428,401 |
| Sequence convolutional | 3,118,409 | 1,573,993 | 747,081 | 990,281 | 1,390,409 |
| Graph convolutional | 3,077,065 | 1,605,993 | 858,953 | 242,793 | 1,381,961 |

**Table S5.**
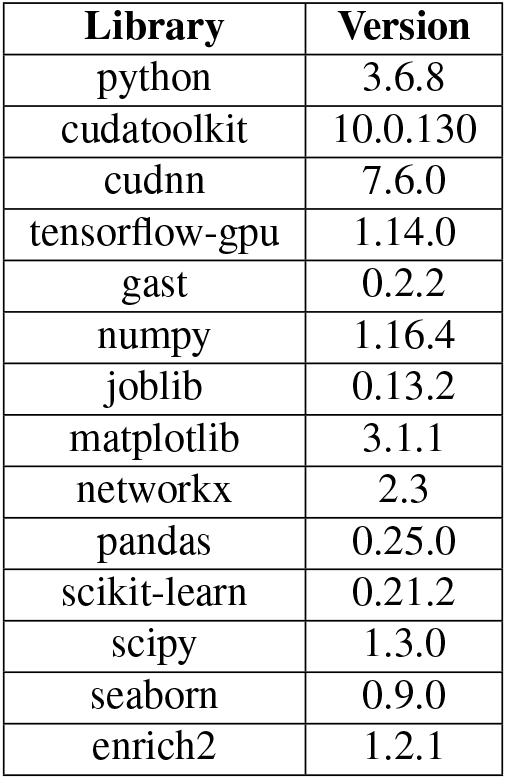
Main software packages. The main libraries and version numbers used to train and evaluate models.

